# The Role of Supraoptic Hypothalamic Arginine Vasopressin Neurons in Aging-Associated Water Balance and Thermoregulatory Deficits

**DOI:** 10.1101/2025.11.03.686336

**Authors:** Nancy Morones, Predrag Jovanovic, Anna Sanetra, Kaitlyn Jang, Nareg Keshishian, Zhihan Cui, Edward Novinbakht, Joshua J. Breunig, Anders Berg, Tamar Pirtskhalava, Selim Chaib, S Ananth Karumanchi, Tamar Tchkonia, Katlin Silm, James L. Kirkland, Celine E. Riera

## Abstract

Aging disrupts physiological homeostasis, impairing thermoregulation, metabolism, and water balance, but the underlying neural mechanisms remain unclear. Here, we identify arginine vasopressin (AVP) neurons in the supraoptic nucleus (SON) of the hypothalamus as a critical driver of these changes. Using single-nucleus RNA-sequencing of the anterior hypothalamus in young and aged mice, we found *Avp* to be one of the most upregulated neuronal transcripts with age. Aged SON^AVP^ neurons displayed enlarged size and heightened excitability, features consistent with hyperactivity. Functionally, chemogenetic activation of SON^AVP^ neurons in young mice reproduced aging-associated phenotypes including hypothermia, reduced energy expenditure, and suppressed water intake. Conversely, knockdown of *Avp* in the SON of aged mice restored water balance, partially improved thermoregulation and systemic metabolism. Pharmacological inhibition of AVP receptors revealed that neuroendocrine release of AVP drives homeostatic deficits, with distinct roles for V1A and V2 receptors. Senolytic drug treatment improved systemic metabolism and reduced inflammaging but does not rescue hypothalamic AVP dysfunction, underscoring a brain autonomous mechanism of age-related physiological failure. Together, our findings establish SON^AVP^ neuronal hyperactivity as a driver of impaired homeostasis with age and suggest that targeted modulation of neuroendocrine AVP signaling may offer a therapeutic strategy to alleviate age-associated water balance defects.

## Introduction

Aging disrupts systemic homeostasis, increasing vulnerability to cardiometabolic and neurodegenerative diseases^1–3^. A hallmark of the aging process is the reduced ability to adapt to physiological challenges, such as maintaining thermoregulation, fluid balance, and metabolic flexibility. While these deficits are well recognized, the underlying neural mechanisms that drive age-associated loss of homeostatic resilience remain poorly defined.

Maintaining body temperature within narrow limits is essential for optimal physiological function and survival. Older adults display blunted responses to both cold and heat stress, increasing risk for hypothermia and hyperthermia^4,5^. With advancing age, both peripheral vasoconstriction and thermogenic heat production are blunted, leading to impaired defense against cold stress^6^. Conversely, during heat stress, diminished sweating, cutaneous vasodilation, and cardiac output increase susceptibility to vital organ damage, including injury to the brain, heart, and kidneys^6,7^. Epidemiological studies underscore the consequences of these deficits, showing that the majority of fatalities from hypothermia or hyperthermia occur in older individuals^8–10^.

Fluid balance is similarly vulnerable to aging. Dehydration is among the ten most frequent diagnoses leading to hospital admission in older adults in the United States^11^. Electrolyte imbalances, including hyponatremia (low sodium) and hypernatremia (high sodium), are also more common in the elderly, and are associated with increased incidence of falls, fractures, osteoporosis, and mortality^11,12^. In addition, elderly individuals have reduced capacity to retain body water, characterized by chronic hypovolemia, or reduced blood volume^13–16^. Despite the clinical significance of these impairments, the mechanisms and specific neural cell types responsible for age-related deficits in thermoregulation and water balance remain unclear. Furthermore, it is not known whether common neural pathways underlie the regulation of both processes.

A critical brain region involved in regulating many physiological functions and behaviors is the hypothalamus, particularly in several anterior nuclei of the hypothalamus (aHYP). Neurons in the aHYP have been implicated in temperature regulation, circadian rhythms, sleep, endocrine function, autonomic nervous system control, thirst and hunger^17–22^. We sought to explore cellular and transcriptional changes occurring within the aHYP possibly driving the physiological changes occurring with aging as well as the functional consequences and potential therapeutic routes to attenuate them. To this end, we employed single-nucleus RNA-sequencing (snRNA-seq) to investigate age-dependent transcriptional dysregulation within the aHYP and aimed to characterize physiological outcomes associated with these changes. Our analysis identified AVP, also known as antidiuretic hormone (ADH), as one of the most upregulated neuronal transcripts with age. Specifically, we found AVP neurons are activated with age in the SON. Remarkably this activation occurred with chronological aging but was not observed in transgenic mice with Alzheimer’s disease-like neurodegeneration. We also show that chemogenetic stimulation of SON^AVP^ neurons is associated with reduced water intake and thermoregulatory deficits resembling those that occur with aging. Remarkably, AVP knockdown within the SON alleviated age-dependent water intake and thermoregulatory deficits.

## Results

### Male aging is associated with a decline in energy expenditure, water intake and thermoregulation

To comprehensively identify metabolic phenotypes of aging in a cohort of C57BL/6 mice, we utilized cohorts of young (3-month-old) and aged (22 to 24-month-old) animals. Body weights in wildtype (WT) C57BL/6 mice increase over time until approximately 24 months of age, then progressively decrease^23,24^. As anticipated, 23-month-aged mice had higher weights than young mice, including increased lean and fat mass (**Fig. 1a**). Measuring whole body basal metabolism by indirect calorimetry revealed key differences in energy homeostasis between young and aged animals (**Fig. 1b-f**). Oxygen consumption (VO_2_), a readout for energy expenditure, was remarkably lower with aging (**Fig. 1b**), in accordance with several reports^23–25^. The respiratory exchange ratio (RER), which measures the daily transition between fuel sources was lower in aged mice, indicating reduced carbohydrate metabolism in aged mice^24^. Locomotor activity was also reduced with aging (**Fig. 1d**). Aged mice consumed less water and food than young counterparts **(Fig. 1e, f**). To assess the onset timeline of these changes, we evaluated the metabolic phenotypes of middle aged (15-month-old) males against young (3-month-old) males and found that many of these traits were already present (**Extended Data Fig. 1. a-e**). This may implicate chronic dehydration as a step in the natural aging process, in addition to other deficits in metabolic health. We also studied the sex effects of these phenotypes in middle aged (15-month-old) and young (5-month-old) female mice, with similar phenotypes emerging in females (**Extended Data Fig. 1 f-j**). Analysis of young and aged male blood samples revealed reduced blood osmolality (**Fig. 1g**). In line with altered water balance, higher plasma levels of AVP were observed with age (**Fig. 1h**).

**Fig. 1.**
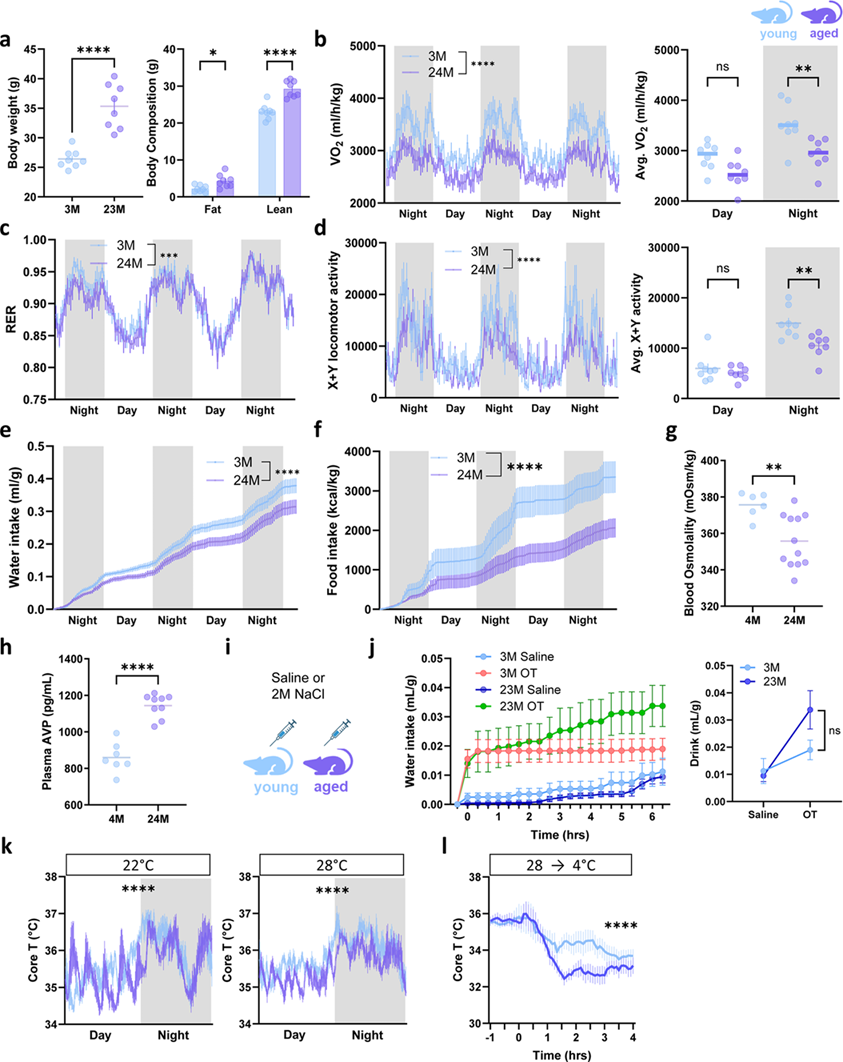
Metabolic and thermoregulatory changes with aging. **a-f**. Metabolic monitoring of male C57BL/6 mice (*n* young = 11, *n* aged = 8) **a.** Body weight (left) and body composition (right). **b-f.** Indirect calorimetry over three days: **b.** VO_2_, **c.** RER, **d.** X+Y locomotor activity, **e.** water intake, **f.** food intake. **g-h.** Postmortem measurements of **g.** Osmolality and **h.** AVP levels in plasma. **i.** Experimental design for inducing osmotic thirst, **j.** water intake plotted over time (left) and by stimulus (right). **k.** Body core temperature (Core T) readings via intraperitoneal Anipill probes over 1 day and 1 night cycle (*n* young = 4, *n* aged = 3) at 22 °C and 28 ° C environmental temperatures. **l.** Core T readings during an acute cold challenge at 4 °C (*n* young = 4, *n* aged = 3). Data are means ± sem. *Student’s t-test (scatterplot, bar graph) or two-way ANOVA (line graph) were used. p-values: ns = p > 0.05; *** = p < 0.005; **** = p < 0.0005*.

AVP is a key hormone involved in the regulation of water balance through the control of water reabsorption from the kidneys^26^. Because of these age-associated differences, we examined the response of aged mice to osmotic thirst using a hypertonic saline infusion test consisting in 2M NaCl injections (**Fig. 1i**), which is a stimulus driving the activation of thirst-sensing neurons in the circumventricular organs (CVO) of the brain^27^. CVO neurons are the primary osmosensitive receptors in the central nervous system (CNS) and modulate the neuronal activity of AVP magnocellular neurons through synaptic connections. Interestingly, despite higher basal AVP levels in aged mice, older animals demonstrated high responsiveness to osmotic thirst (**Fig. 1j**), implying that the ability to respond to a thirst stimulus detected by hypothalamic osmoreceptors is not impaired during aging^28^. To study thermoregulatory differences with age, we subjected mice to various ambient temperatures, including mild cold (22°C), thermoneutrality (28°C) and acute cold (4°C) and measured their core temperature (Core T) using Anipill capsules implanted in the peritoneal cavity (**Fig. 1k,l**). Continuous Core T monitoring corroborated thermoregulatory deficits in aged mice maintained at 22°C and 28°C housing temperatures (**Fig. 1k**). In both conditions, Core T in the aged group appeared to be lower than in young mice, particularly during the light phase. To evaluate cold defenses with aging, we exposed mice to an acute 4°C ambient temperature and observed a more rapid decline in Core T in the aged mice compared to young mice (**Fig. 1l**), indicative of impaired thermoregulation with age upon extreme cold exposure^29^.

### Molecular and cellular features of aging in the male anterior hypothalamus using snRNAseq identifies increased transcriptional Avp

After investigating age-related physiological variations in energy balance, water consumption, thermoregulation, and physical activity, our goal was to pinpoint the cellular factors accountable for these characteristics within the brain. The anterior region of the aHYP contains key nuclei including the preoptic area (POA), paraventricular nucleus (PVN), suprachiasmatic nucleus (SCN), ventromedial (VMH) and SON hypothalamic regions which are involved in thermoregulation, circadian rhythms, energy balance, neuroendocrine regulation, thirst and hunger^17–22^. To investigate cellular changes associated with aging in the aHYP, we micro-dissected this region in mice and isolated nuclei from young and aged sample pools for single-nucleus RNA-sequencing (snRNA-seq) (**Fig. 2a**). After alignment and quality control, we visualized 18,287 nuclei (10,267 young; 8020 aged) by Uniform Manifold Approximation and Projection (UMAP) using Seurat (**Fig 2b**) and annotated clusters into cell types based on canonical markers (**Fig. 2c, Extended Data Fig. 2a**)^27,30^. Interestingly, we saw no major differences in UMAP projections between our young and aged sample pools at this level (**Fig. 2d**). However, examination of cell-type population proportions showed a decrease in astrocytes in aged mice (**Fig. 2e**), which has been previously reported^31,32^. We also noted a proportional increase of oligodendrocytes (**Fig. 2e**), which has been characterized in neurodegenerative diseases such as Alzheimer’s disease (AD) and Lewy body dementia^33^. Further analysis of glial cell types by RNA velocity in Scanpy (**Extended Data Fig. 3a-b**) revealed that microglial populations exhibit distinct clustering between young and aged mice (**Extended Data Fig. 3c**). Cell directionality and velocity pseudotime further convey aging-related differences (**Extended Data Fig. 3d-e**). However, there was not a clear correlation between RNA velocity driver genes and cell clusters (**Extended Data Fig. 3f-i**).

**Fig. 2.**
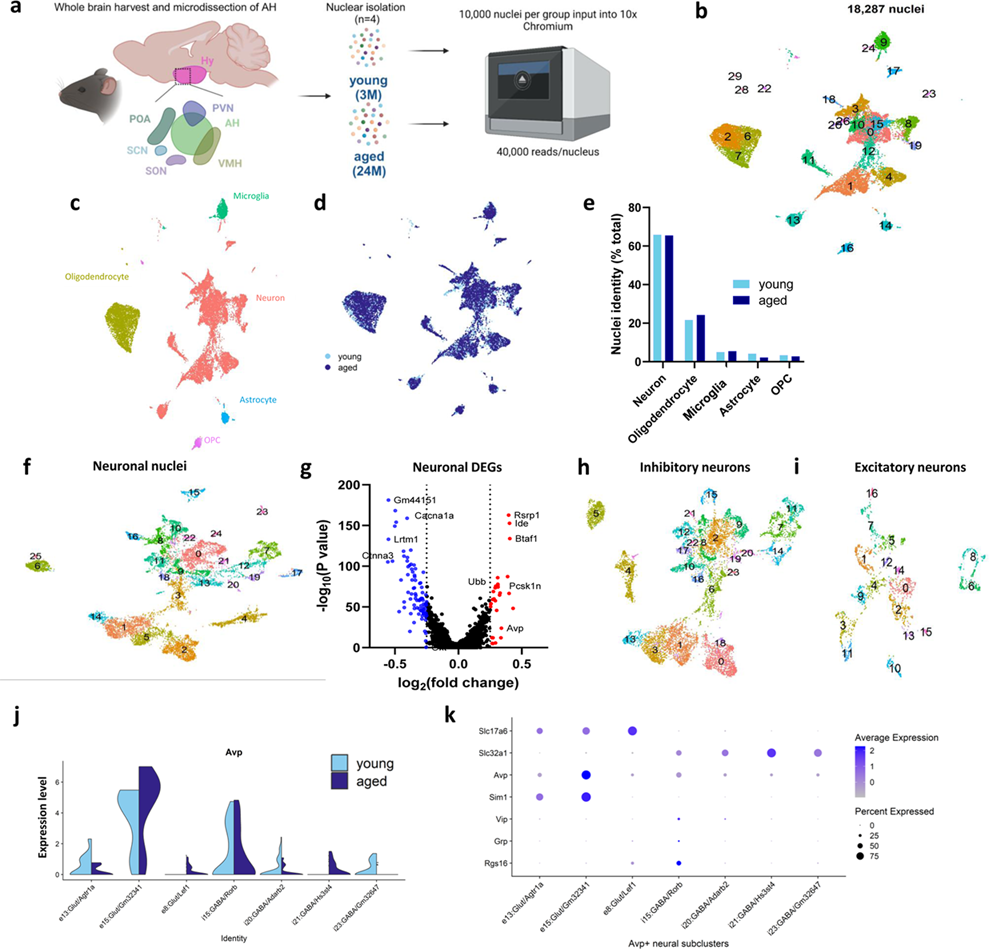
Single-nuclei RNA sequencing of the aging anterior hypothalamus. **a.** Schematic illustrating workflow of microdissection and age groups pooled into snRNA-seq (*n*=4). **b.** UMAP of all nuclei from both experimental groups, processed into 29 clusters. **c.** UMAP of 5 major cell types identified in dataset. **d.** UMAP of subset neural clusters, identifying 25 subclusters. **e.** Cell proportions across each cell type split by age. **f. g.** Volcano plot of differentially expressed genes in aging vs. young nuclei pools. **h-i.** UMAP of inhibitory and excitatory neurons with respective clusters. **j.** Violin plot of Avp expression across Avp-rich clusters from excitatory (e13, e15, e8) and inhibitory (i15, i20, i21, i23) subclusters, with increased Avp expression observed in e15. **k.** Dot plot of key gene markers in Avp-rich clusters including Sim1 (marker of PVN and SON neurons) and Rgs16 (labels SCN neurons) shows that high e15 expression of Avp coincides with Sim1 expression.

Because of their role in physiology, we chose to focus on the neural population and reclustered it into 25 new clusters (**Fig. 2f**). From these subset cells, we performed differential gene expression analysis (DEG) to identify transcriptional changes in neurons with aging (**Fig. 2g**). Among these, *Avp* was identified within the top enriched transcripts of aged neuronal populations. To further elucidate the identity of the *Avp*-enriched neurons, we selected subset clusters that were enriched in either GABAergic *Slc32a1* or glutamatergic *Slc17a6*, markers for inhibitory and excitatory neurons^34^, respectively, and re-clustered them into UMAP (**Extended Data Fig. 2b-d**). Queries of *Avp* across these subclusters showed enrichment in 3 excitatory clusters and 4 inhibitory clusters (**Fig. 2j**). Among these, excitatory cluster e15:Glut/Gm32341 exhibited *Avp* enrichment even by unbiased FindAllMarkers query (**Extended Data Fig. 2d**). To unmask potential localization of these clusters, we surveyed known markers of *Avp*-enriched regions^35^ including the PVN, SCN and SON (**Fig. 2k**). The e15 cluster containing *Avp* coincided with *Sim1*, a cellular marker labelling multiple categories of PVN and SON neurons^35^, but not with *Rgs16* which is more highly associated with SCN neurons^35,36^. These data suggest that selective subregions, including PVN and SON expressing *Avp* might become enriched with aging.

### Characterization of age-dependent changes in hypothalamic AVP neuron cellular composition reveals neural soma enlargement in the SON

Having found upregulation of *Avp* in excitatory neurons with age, we investigated potential age-related cellular changes within the hypothalamic neuronal populations that produce AVP. We tested if there is a link between transcription and translation of AVP using immunohistochemistry of mouse hypothalamic sections with a vasopressin antibody, targeting the AVP neuropeptide. We analyzed various cellular features of AVP neurons within the PVN, SCN, and SON regions, which were the exclusive regions of the aHYP that produce this transcript according to the snRNAseq data (**Fig. 3a-c**). The comparison of young (3-month-old) and aged WT (24-month-old) samples indicated that the cell size of AVP neurons was enlarged in the aging SON compared with young controls (**Fig. 3e**). We found no significant differences in individual region size (**Fig. 3f**), AVP-positive neuronal count (**Fig. 3g**), or AVP signal intensity per region (**Fig. 3h**). Notably, there was no significant difference in cellular roundness or AVP content per cell, measured using fluorescence intensity between ages for each region (**Fig. 3i, j**). Interestingly, changes in AVP neuronal size have been correlated with increased neuronal activity and peptidergic production^37–40^, therefore indicating that increased transcriptional expression of *Avp* observed by snRNAseq leads to higher production of AVP with SON^AVP^ neurons. These findings are in accord with human aging data, where post-mortem analysis of human brains has revealed an increase in the size of AVP neuronal somas, nucleoli, and the Golgi apparatus, in older patients (80-100 years-old)^37–39,41^.

**Fig. 3.**
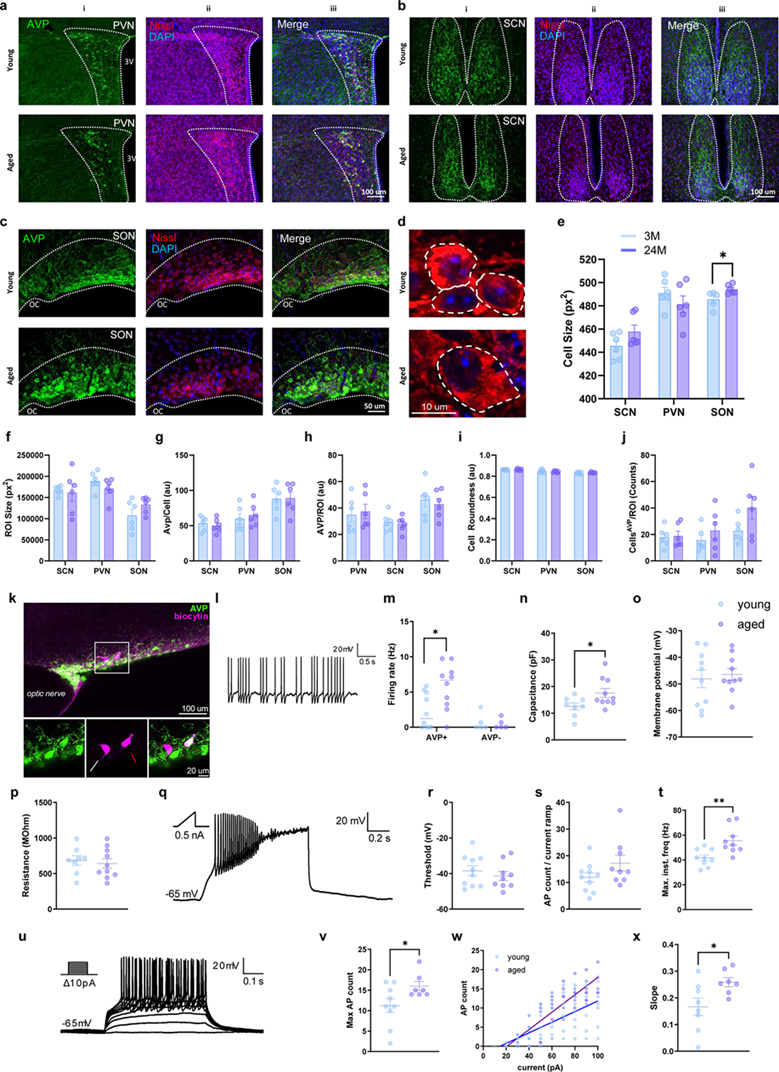
Analysis of AVP expressing cell bodies in PVN, SCN, and SON. **a-c.** Representative immunohistochemistry image of AVP^+^ cell bodies in young and old male mice, (i) anti-AVP (green), (ii) Nissl (magenta) and DAPI (blue), iii Merge image. **a.** PVN, **b.** SCN, **c.** SON. **d.** High magnification images of AVP^+^ neurons in young and old mice, AVP (red) and DAPI (blue). **e-j.** Quantification of morphological features of AVP neurons in each region (*n* young = 5, *n* aged = 6). **e.** AVP cell body size, **f.** ROI size, **g.** AVP cell bodies per ROI, **h.** AVP signal intensity per ROI, **i.** AVP cell roundness, and **j.** AVP signal intensity per cell. **k.** Representative image of AVP^+^ (red arrow) and AVP^-^ (white arrow) neurons recorded from slice electrophysiology, anti-AVP (green), biocytin (magenta). **l.** Representative trace of a baseline recording at 0 A holding current from AVP^+^ neuron. **m.** Spontaneous firing rate of AVP^+^, but not AVP-negative cells (p>0.9999, Mann Whitney test). **n.** Comparison of membrane capacitance between young and old AVP^+^ neurons. **o.** Resting membrane potential. **q.** A 1s, 0.5 nA current ramp was applied from a manually adjusted membrane potential to −65 mV to determine firing threshold and highest possible firing ability. **r.** Threshold, **s.** total action potential (AP) count. **t.** maximum instantaneous frequency. **u.** With membrane voltage still adjusted to −65 mv, 10 depolarizing current steps (from 0 to 100 pA, 10 pA increments) were applied to establish the gain in firing rate over increasing stimulation strength for **v.** maximum AP count, **w.** gain value defined as the slope of a linear regression fit to the number of evoked APs at successive steps. **x.** Slope values of w. Data are means ± sem*. One-way ANOVA or* S*tudent’s t-test were used. p-values*: **\*** = p < 0.05.

### Ex vivo electrophysiology supports higher excitability with age in SON^AVP^ neurons

To assess changes in neuronal excitability, we employed slice patch clamp recording of hypothalamic neurons. We recorded a total of 30 neurons, 20 of which were AVP+ (10 from young and 10 from old animals) and 10 AVP-(5 per group), as revealed by post-recording immunostaining (**Fig. 3k-m**). When recording spontaneous activity in current clamp mode (0 A holding current, **Fig. 3l**), AVP+ cells from old animals showed an increase in firing rate, which was significantly different from the AVP+ cells from young animals **(Fig. 3m**). On the other hand, the firing rate of AVP-neurons did not differ significantly between the age groups. Consistent with the observation of a larger soma in old SON^AVP^ neurons, we observed increased membrane capacitance in these cells (**Fig. 3n**). However, neither resting membrane potential (**Fig. 3o**) nor membrane resistance was different between the groups (**Fig. 3p**), excluding increased leak current as a potential factor underlying the increased firing rate. To measure excitability we first manually brought each cell down below their firing threshold (−65 mV) and then applied a 1-s long, 0.5 nA current ramp to measure the voltage at which the cells fire the first action potential (AP) (threshold), the number of APs evoked during the ramp before reaching depolarization block, and the highest possible firing rate, or maximum instantaneous frequency (**Fig. 3q-t**). While the AP firing threshold (**Fig. 3r**) and the total AP count during the ramp (**Fig. 3s**) were not significantly different between the age groups, cells recorded from old animals reached a higher maximal frequency (**Fig. 3t**), indicating that these cells are more excitable. Therefore, as the last step, while still holding the cells at −65 mV, we applied a series of 0.5s long current steps ranging from 10 to 100 pA (10 pA increments), to determine the maximum number of APs evoked during stimulation at a constant magnitude without reaching supraphysiological levels. We also calculated the gain of the firing rate at successive pulses, defined here as the slope of the linear function fitted to the relationship between the pulse amplitude and the number of APs evoked (**Fig. 3u-x**). Both parameters were increased in the old group, supporting increased excitability of the aged neurons, especially in response to stronger stimuli. Overall, these results indicate that SON^AVP^ neurons become more excitable with age, already firing at a higher rate at baseline but also responding more strongly to a depolarizing input.

### Distinct hypothalamic and homeostatic phenotypes in chronological aging and Alzheimer’s disease like pathology in mice

Having established selective neuronal dysfunction of the SON with chronological aging, we investigated this system in a model of age-dependent neurodegeneration, representative of AD β-amyloid accumulation. Disruptions in circadian rhythm, including thermoregulatory and sleep-wake cycles changes, are commonly reported in AD patients^42–44^. As AD severity increases, these circadian disturbances become more pronounced^45,46^. To evaluate whether the cellular deficits observed with chronological aging are mirrored in age-dependent neurogenerative disease, we employed familial AD mice (5xFAD), which overexpress 5 human mutations, 3 in the amyloid-beta precursor protein (APP) and 2 in the presenilin-1 (PS1) genes^47^. We used 6-month-old 5xFAD mice compared to WT littermate controls, as this model has early onset neurodegeneration^48^. While metabolic traits differed between the 5xFAD and controls, these changes were inverse to those found with chronological aging (**Extended Data Fig. 4**). As previously described^49^, 5xFAD mice had reduced adiposity and increased energy expenditure **(Extended Data Fig. 4a-c**), no significant difference in food intake (**Extended Data Fig. 4d**), minor changes in activity (**Extended Data Fig. 4e**) and higher water intake than WT, as opposed to chronological aging (**Extended Data Fig. 4f**). Their Core T was higher during the night cycle at 22°C and 28°C ambient temperatures (**Extended Data Fig. 4g,h**). Remarkably, their cold defense mechanisms were largely better compared to controls in response to extreme cold challenge (**Extended Data Fig. 4i**). Together, these results indicate that key metabolic differences associated with normal aging are not replicated in the neurodegenerative 5xFAD mouse model.

When analyzing the cellular features of hypothalamic AVP neurons in 5xFAD mice compared to WT, an imbalance in the AVP system was observed but differed from the features of chronological aging. The SON region was enlarged in AD mice, concurrent with an increased number of AVP neurons and a paradoxical reduction in AVP signal intensity in the region **(Extended Data Fig. 4j-m)**. However, cell size of SON neurons in AD mice was not significantly increased (**Extended Data Fig. 4p**). Taken together, these findings highlight key differences in the AVP system between chronological aging and murine AD-like pathology, with marked differences observed in the SON.

### Chemogenetic stimulation of AVP neurons in the SON leads to hypothermia, hypometabolism and reduced water uptake

The specific roles of centrally released AVP from the various hypothalamic regions in metabolism have not been resolved. Previously, chemogenetic stimulation of all AVP neurons using a genetic rat line expressing the human muscarinic acetylcholine receptor (hM3Dq) led to reduced feeding, drinking and urine volume^50^, but the contribution of the more specific AVP hypothalamic nuclei and associated subpopulations to these responses, as well as their role in thermoregulation, remains unclear. Interestingly, activation of PVN^AVP^ neurons also acutely suppressed food intake in mice^51^.

To investigate the role of SON^AVP^ neurons in energy homeostasis, thermoregulation, and water intake, we interrogated their function using viral chemogenetics^52^ and simultaneous recording of core T using Anipill probe implants. We delivered AAV8-DIO-hM3D-mCherry bilaterally in the SON of AVP-IRES-Cre male mice and confirmed accurate targeting of the region (**Fig. 4a, b**). Circulating AVP levels fluctuate upon stimulation of magnocellular neurons of the SON, responsible for synthesizing AVP, which is then deposited to the posterior pituitary and secreted into the bloodstream. Acute stimulation of these neurons using intraperitoneal injection of their ligand deschloroclozapine (DCZ) at 1mg/kg resulted in increased plasma AVP as measured by Copeptin, a stable AVP precursor^53–55^, as well as lower plasma osmolality (**Fig. 4c**). Remarkably, DCZ injection in AVP-M3D mice led to transient hypothermia, the magnitude of which was increased with high dose (1mg/kg) over low dose DCZ (0.1mg/kg) in comparison to saline treatment (**Fig. 4d**). Similarly, we observed DCZ dose-dependent impairment in energy expenditure as characterized by lower VO_2_ and concomitant reduced locomotor activity, as well as suppressed water intake without significant effect on food intake (**Fig. 4e-h**). Water intake was suppressed for over 4 hours following DCZ injection (**Fig. 4h**).

**Fig. 4.**
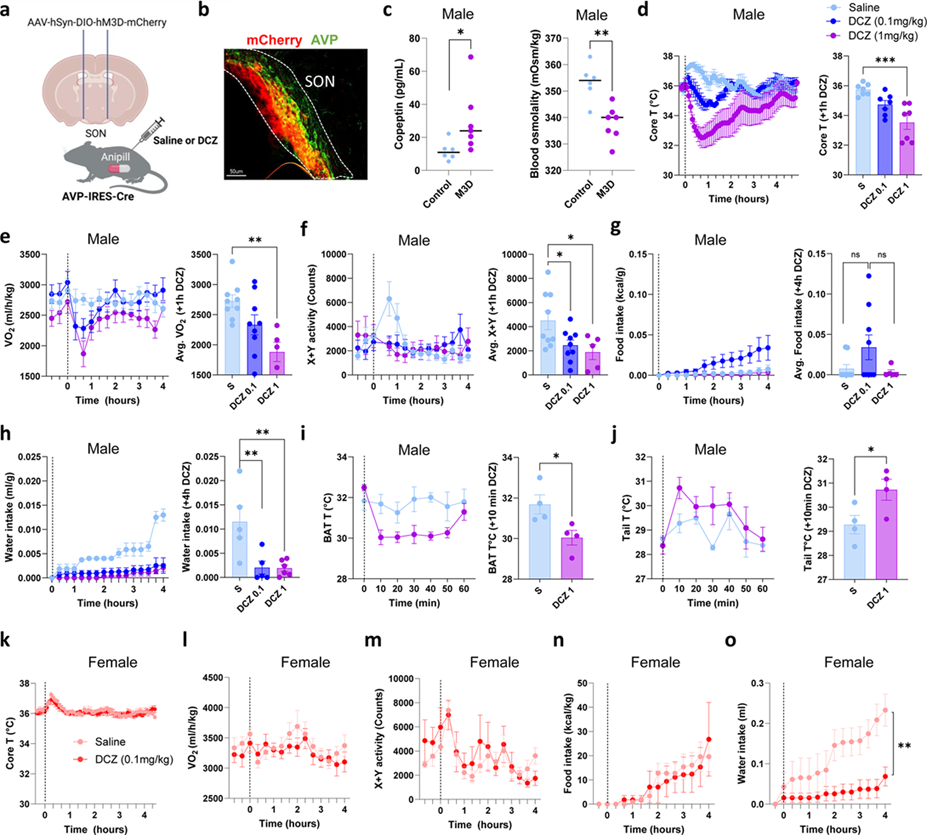
Chemogenetic stimulation of SON^AVP^ neurons. **a.** Experimental Design: AVP-IRES-Cre mice received bilateral stereotaxic injections of AAV-hSyn-DIO-hM3D-mCherry followed by intraperitoneal (I.P.) delivery of saline or DCZ in male **(c-j)** and female mice **(k-o). b.** Immunohistochemistry of mCherry (red) and AVP (green) overlapping in SON. **c.** Plasma levels of copeptin, and blood osmolality between AVP-M3D mice (n=5) and saline controls (n=6) post-injection of DCZ. **d.** Core T plotted overtime (left) and averaged over 1^st^ hour post-injection (right) of AVP-M3D mice receiving either saline, DCZ (0.1mg.kg) or DCZ (1mg/kg). **e-h.** Indirect calorimetry assessment of AVP-M3D mice receiving either saline, DCZ (0.1mg.kg) or DCZ (1mg/kg): **(e)** VO_2_, **(f)** X+Y locomotor activity, **(g)** food intake and (**h)** water intake. **i-j.** Infrared (IR) temperature readings of **(i)** brown adipose tissue (BAT T) and **(j)** tail (Tail T). **k-o.** Female AVP-M3D mice receiving either saline or DCZ (0.1mg.kg): (**k)** Core T, (**l)** VO_2_, (**m)** locomotor activity, (**n)** food intake, and (**o)** water intake. Data are means ± sem. *One-way ANOVA or Student’s t-test (bar graphs), two-way ANOVA (line-graphs) were used. p-values: **\*** = p < 0.05*.

To investigate the physiological role of SON^AVP^ neurons in thermoregulation, we assessed changes in skin temperatures using infrared imaging in the brown adipose tissue (BAT) and tail base areas. DCZ induced an acute BAT temperature decrease, which indicates suppression of BAT thermogenesis, as well as increased tail temperature, indicating vasodilation leading to heat loss (**Fig. 4i,j**). Both measurements are consistent with prior reports of mouse hypothermia induced by decreased BAT thermogenesis combined with enhanced tail vasodilation^56^. These data also highly suggest that the drop in energy expenditure observed upon stimulation of SON^AVP^ neurons is mediated by rapid inhibition of BAT thermogenesis, vasodilation of blood vessels and concurrent hypothermia.

We explored whether female SON^AVP^ neurons were linked to similar metabolic phenotypes to those in males. To our surprise, we found that acute DCZ administration did not produce significant hypothermia in female mice (**Fig. 4k**), nor did it reduce energy expenditure, locomotor activity, or feeding (**Fig. 4l-n**). However, as in males, DCZ injection dramatically suppressed water intake (**Fig. 4o**).

### Pharmacological inhibition of vasopressin receptors uncouples water intake and thermoregulatory output linked to activation of SON-AVP neurons

AVP synthesis occurs predominantly in the SON and in small quantities within the PVN^57^. Storage vesicles containing AVP are transported down the axons through the hypothalamic-hypophysial tract, where they are ultimately released in the posterior pituitary. The secreted peptides then enter the body’s systemic circulation. Additionally, central AVP acts as a neurotransmitter or neuromodulator, and has sites of action on autonomic brainstem centers^58^. AVP administered intraventricularly in rodent models causes sympathetic activation^59^, and therefore could account for catecholamine (such as noradrenaline) release and modulation of vasodilation and thermogenesis through adipose adrenergic receptors. To investigate whether the metabolic effects associated with SON neuronal activation depend on circulating or central AVP action, we acutely injected wild-type mice implanted with Anipill probes with saline or AVP peptide (0.1mg/kg) intraperitoneal injections (**Extended Data Fig. 5a**). Similarly to the chemogenetic stimulation of SON^AVP^ neurons, AVP peptide injection induced a hypothermic response that was resolved shortly after 1 hour (**Extended Data Fig. 5b,c**). BAT skin temperature was concomitantly reduced and tail temperature also dropped (**Extended Data Fig. 5d,e**). AVP is a major vasoconstrictor of vascular smooth muscle cells, in line with our intraperitoneal peptide injection on tail temperature, but has been shown to also have vasodilator capacity depending on the regional site of action^60^. These data suggest that central SON^AVP^ stimulation (**Fig. 4j**) has diverging or additional effects on blood vessel constriction compared to peripheral AVP (**Extended Data Fig. 5e**).

We then aimed to identify potential target receptors associated with peripheral AVP action on metabolism. We employed vasopressin receptor antagonists (10mg/kg I.P.) that inhibit peripheral vasopressin action^61^: the V1AR antagonist SR49059 and FDA-approved V2R antagonist Tolvaptan. These drugs were injected 20 min prior to AVP injection at the onset of the night cycle (**Fig. 5a**). We found that AVP induced a significant decrease in VO_2_, which was suppressed by V1AR inhibition (**Fig. 5 b, c**). AVP peptide injection also powerfully suppressed water intake for up to 3 hours, and this phenotype was largely reversed by V2R inhibition (**Fig. 5d, e**) without significantly affecting food intake (**Fig. 5f**). V1AR inhibition also restored water intake upon AVP injection (**Fig. 5d, e**). Similarly, hypothermia induced by AVP was reversed by V1AR inhibition (**Fig. 5g, h**).

**Fig. 5.**
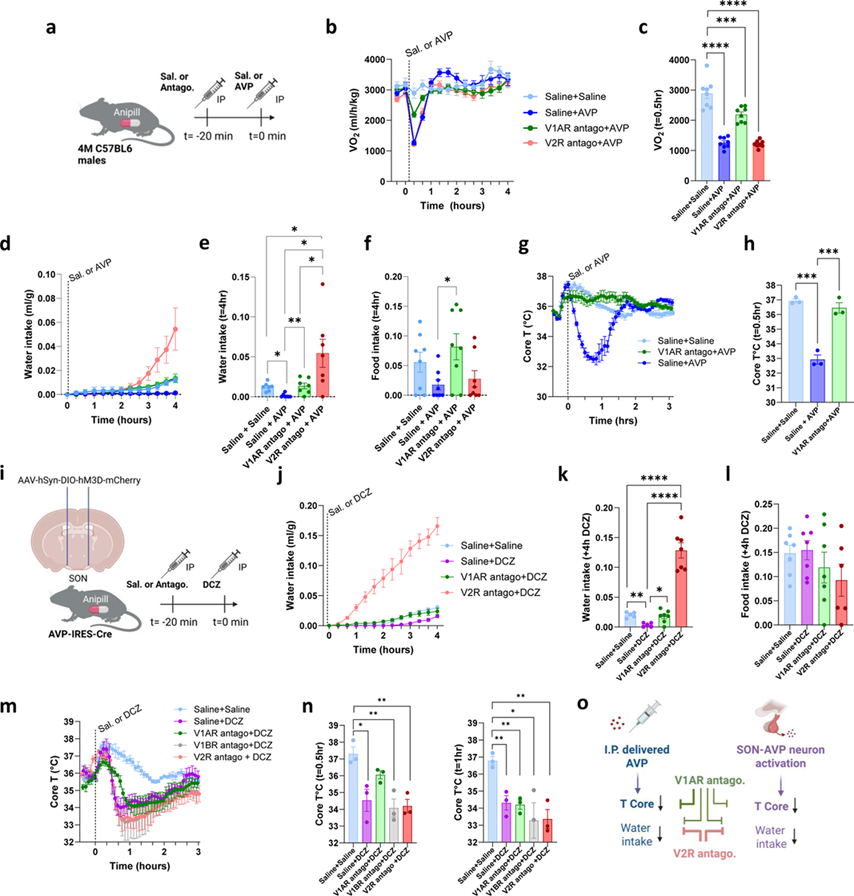
Pharmacological targeting of AVP receptors in AVP peptide-treated and SON chemogenetic modulation of AVP^+^ neurons. **a.** Experimental Design for (b-h): C57BL6/J male mice implanted with Anipill temperature probes in the abdominal cavity were I.P. injected with saline (Sal.) or antagonist (Antago.) at t=-20 min followed by either I.P. saline or 0.1mg/kg AVP peptide at t=0 min (*n = 7-8*). **B-c.** VO_2_ over time and (c) at t=0.5 hr following injection. **d-e.** Water intake over time and (**e)** at t=4 hrs following injection. **f.** Food intake at t=4 hrs following injection. **g-h.** Core T over time and (**h)** at t=0.5 hr following injection. **i.** Experimental Design for (**j-n**): male AVP-M3D mice with Anipill implants were I.P. injected with saline (Sal.) or antagonist (Antago.) at t=-20 min followed by either I.P. saline or 1mg/kg DCZ at t=0 min (*n = 5-7*). **J-k.** Water intake over 4 hours following DCZ delivery and **(k)** at t=4 hrs following injection. **l.** Food intake post-DCZ treatment at t=4 hrs following injection. **m-n.** Core T over time and at (**n**) t=0.5hr (left), t=1hr (right). **o.** Schematic summary demonstrating similar effects of I.P. delivered AVP and SON^AVP^ neuron activation on vasopressin receptors in vivo. Data are means ± sem. *One-way ANOVA. p-values: ns = p > 0.05; *** = p < 0.005; **** = p < 0.0005*.

We then employed these antagonists to treat AVP-M3D mice receiving saline or DCZ (1mg/kg) (**Fig. 5i**). Mirroring AVP peptide injections, we found that V2R inhibition enhanced water intake in DCZ-treated mice, and V1AR inhibition restored water uptake back to control levels (**Fig. 5j,k**). These effects were independent of food intake changes (**Fig. 5l**). We included the V1BR antagonist TASP0390325 which mediates the vasopressin dependent release of ACTH in pituitary corticotrophs^62^, to determine if this pathway was recruited in the hypothermic response induced by SON^AVP^ neuronal stimulation. As with AVP peptide injections, we found that V1AR inhibition transiently suppressed DCZ-induced hypothermia, but the effects of the V1AR antagonist were only notable for 30 min following DCZ administration and not after 1 hour (**Fig. 5 m,n**), suggesting that the chemogenetic stimulation of neurons outlasted the presence of the antagonist in circulation. V2R and V1BR inhibition had no effects on AVP-M3D dependent hypothermia. Taken together, these findings demonstrate that hypothalamic recruitment of AVP neurons depends on the neuroendocrine release of AVP, which regulates core temperature by V1AR and modulates water intake through the combined actions of V1AR and V2R (**Fig. 5o**).

Following this, we postulated that aging would more closely resemble a hyperactive AVP neuronal state and thus, pharmacological inhibition would have similar therapeutic benefits to those in our chemogenetic model. We tested this by treating mice with acute injections of V1AR and V2R antagonists and measuring potential improvements in age-associated metabolic phenotypes (**Extended Data Fig. 5f**). Energy expenditure (VO_2_), RER, and locomotor activity were not impacted significantly by acute antagonist treatment (**Extended Data Fig. 5g-i**). Water intake was largely enhanced by V2R antagonist treatment, and more modestly by V1AR antagonist treatment (**Extended Data Fig. 5j,k**) without significantly impacting food intake (**Extended Data Fig. 5l**). No other notable metabolic changes were observed upon acute injection of these drugs, including thermoregulation at ambient and cold temperatures (**Extended Data Fig. 5 m,n**). This suggests that the water balance system exhibits greater plasticity than the thermoregulatory system, whose age-related impairments cannot be readily reversed through acute interventions.

### Downregulating AVP in aged mice enhances water intake and systemic metabolic resilience

To explore the therapeutic potential of reduced *Avp* transcription in aging and metabolism, we targeted SON^AVP^ neurons using a lentiviral packaged shRNA in the aging brain. AVP-shRNA or control non-targeting mCMV-TurboGFP lentivirus were injected in the SON of 22-month-old male mice (**Fig. 6a**). Efficient knockdown of *Avp* within the SON was confirmed using immunohistochemistry (**Extended Data Fig. 6a**) and RNAScope (**Fig. 6b-c; Extended Data Fig. 6b,c**). Knockdown mice had decreased circulating levels of AVP (**Fig. 6d**) and a trend towards higher blood osmolality (**Fig. 6e**). Animals were monitored over a period of 6 weeks, with their weights and body composition unchanged over time (**Fig. 6, Extended Data Fig. 6d**). Indirect calorimetry revealed no significant difference in oxygen consumption between control and *Avp* shRNA animals (**Fig. 6h**). Locomotor activity was also unchanged (**Fig 6i**). However, the RER was more elevated upon *Avp* knockdown (**Fig. 6j**), suggesting that substrate utilization was shifted towards increased carbohydrate metabolism by *Avp* knockdown, a metabolic trait that is present in younger animals (**Fig. 1c**)^24^. Interestingly, *Avp* knockdown in the SON also increased water intake (**Fig. 6k**), towards more youthful drinking levels without significantly affecting food intake (**Fig 6l**). The analysis of thermoregulation in these mice revealed that *Avp* knockdown was associated with a slight trend towards warmer Core T during light and dark cycles at 22°C (**Fig. 6m**), but not at 28°C (**Fig. 6n**). *Avp* knockdown was characterized by a tendency to a shorter hypothermic phase at 4°C compared to aged-matched controls, albeit not statistically significant (**Fig. 6o**). These data support the notion that transcriptional suppression of AVP in SON improves water intake, enhances carbohydrate metabolism and has mild benefits on thermoregulation in mouse aging.

**Fig. 6.**
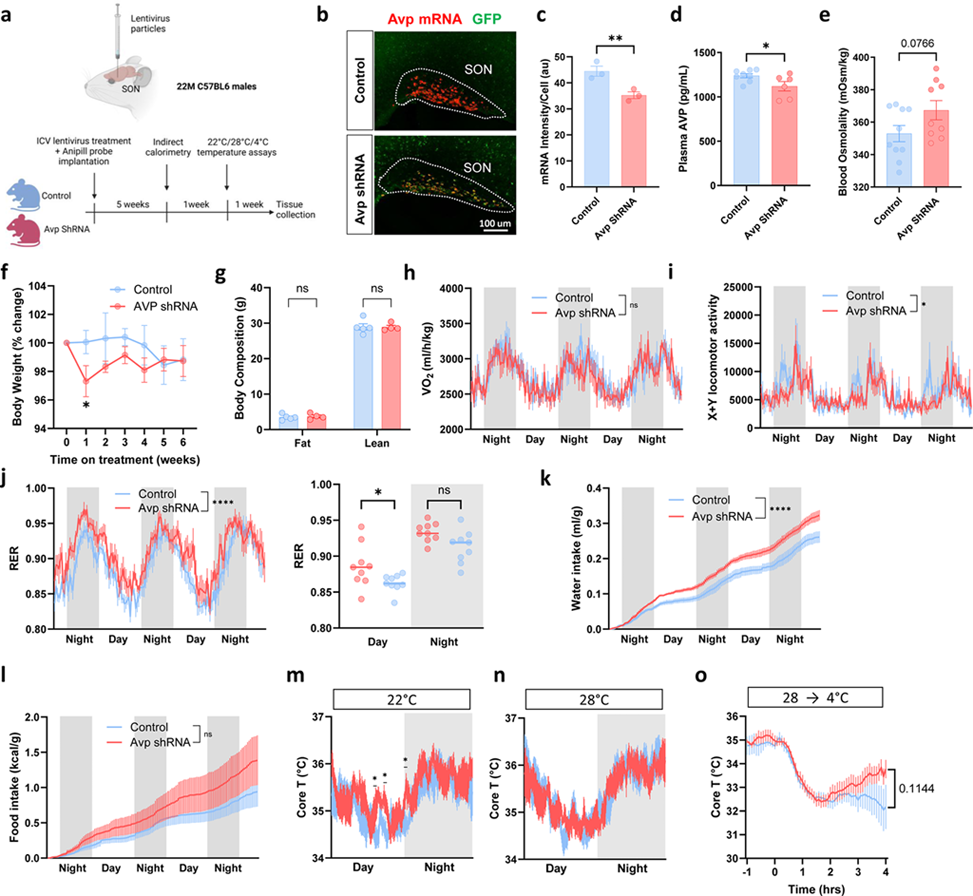
*Avp* silencing in the SON increases water intake and improves energy utilization in aged mice. **a.** Schematic of the experiment**. b**. Representative RNAScope image of Avp-positive cell bodies in the SON (green, GFP Viral particles; red, Avp mRNA). **c.** Quantification of mRNA intensity per cell (*n* = *3*). **d.** Plasma AVP levels measured at sacrifice (*n* = *6-8*). **e.** Blood osmolality measured at sacrifice (*n* = *9-10*). **f.** Percent change in body weights of male mice during treatment (*n* = *9-10*). **g.** Fat and lean mass at sacrifice (*n* = *5*). **h-l.** Indirect calorimetry assessment of VO_2_, locomotion, RER, water and food intake across mice (*n = 9*). **m,n.** Core T measurements using intraperitoneal Anipill probes over 1 day and 1 night cycle (*n* = 7) at **(m)** 22°C and **(n)** 28 °C environmental temperatures. **o.** Core T readings from 4°C cold challenge (*n* = 7). Data are means ± sem. *Student’s t-test (bar graphs) or Two-way ANOVA (line graphs) were used. p-values: ns = p > 0.05; *** = p < 0.005; **** = p < 0.0005*.

### Senolytic drugs improve systemic metabolism but do not significantly restore water balance and thermoregulatory defects with age

Senescent cells, which are in essentially permanent cell cycle arrest, can accumulate with age in many organs in both experimental animals and humans^63^. Many of these cells develop a senescence-associated secretory phenotype (SASP), which consists of pro-inflammatory, pro-apoptotic and pro-fibrotic factors, among others^63^. Accumulation of such SASP secreting cells is associated with a chronic low-grade inflammatory state associated with aging and chronic diseases, known as inflammaging. The senolytic cocktail Dasatinib and Quercetin (D+Q) can eliminate these cells and improve metabolic function, as well as systemic inflammation^64,65^. In addition, D+Q treatment has been shown to relieve neuroinflammation associated with senescent glial cells in the brain of obese mice^66^. To test the role of inflammaging in triggering the abnormal neuronal activation of SON^AVP^ neurons, we investigated the outcome of targeting senescent cells using D+Q on vasopressin-related physiological decline with age. Because biological aging phenotypes can be heterogenous we pre-screened our mice and grouped them into evenly distributed weights and metabolic phenotypes (**Extended Data Fig. 7a-e**).

We treated aged and young mice with D+Q or Sham gavage (**Fig. 7a**). D+Q treatment did not alter weight significantly over seven weeks of repeated treatment (**Fig. 7b**), nor did it impact overall appearance (**Fig. 7c**). The efficacy of D+Q treatment was validated in gonadal WAT, with a trend towards reduction in p16 expression and tumor necrosis factor-alpha or Tnfa (**Fig. 7d**). Metabolic phenotypes of aging were present in aged cohorts (>22M) over young counterparts (4M), including weight loss over time (**Fig. 7b**), reduced VO_2_, lower RER, lower spontaneous activity, reduced drinking and feeding compared to young cohorts (**Fig. 7e-i**). VO_2_, RER, and locomotor activity were altered by D+Q in aged cohorts towards more youthful profiles (**Fig. 7e-g, Extended Fig. 7a-e**). However, water and food intake were largely unchanged (**Fig. 7h-i**). In accordance with water intake measurements, blood osmolality was not significantly improved upon D+Q in aged cohorts (**Fig. 7j**). Investigation of thermoregulatory function upon D+Q treatment did not show significant differences in Core T during light and dark cycles at 22°C, 28°C and 4°C (**Fig. 7k,l**). Furthermore, qualitative analysis of SON^AVP^ neurons further showed no notable changes to neural soma in aged mice receiving D+Q compared to Sham gavage (**Fig. 7m-q**). Despite this, we investigated whether D+Q could alleviate neuroinflammation within the SON with age, using immunostaining of cytokine IL-6 and astrocyte reactivity marker glial fibrillary acidic protein (GFAP) (**Extended Fig. 7f-h**). We found that IL-6 levels in the SON remained high in aged D+Q or Sham animals, indicating that high inflammation measured within the aging hypothalamus is not alleviated by oral D+Q treatment in aged mice. These results indicate that D+Q, while providing a crucial metabolic improvement through systemic benefits, did not significantly attenuate hypothalamic neuroinflammation or age-dependent defects in SON^AVP^ neurocircuitry when delivered orally to mice over 22-month-old at doses sufficient to alleviate systemic metabolic dysfunction.

**Fig. 7.**
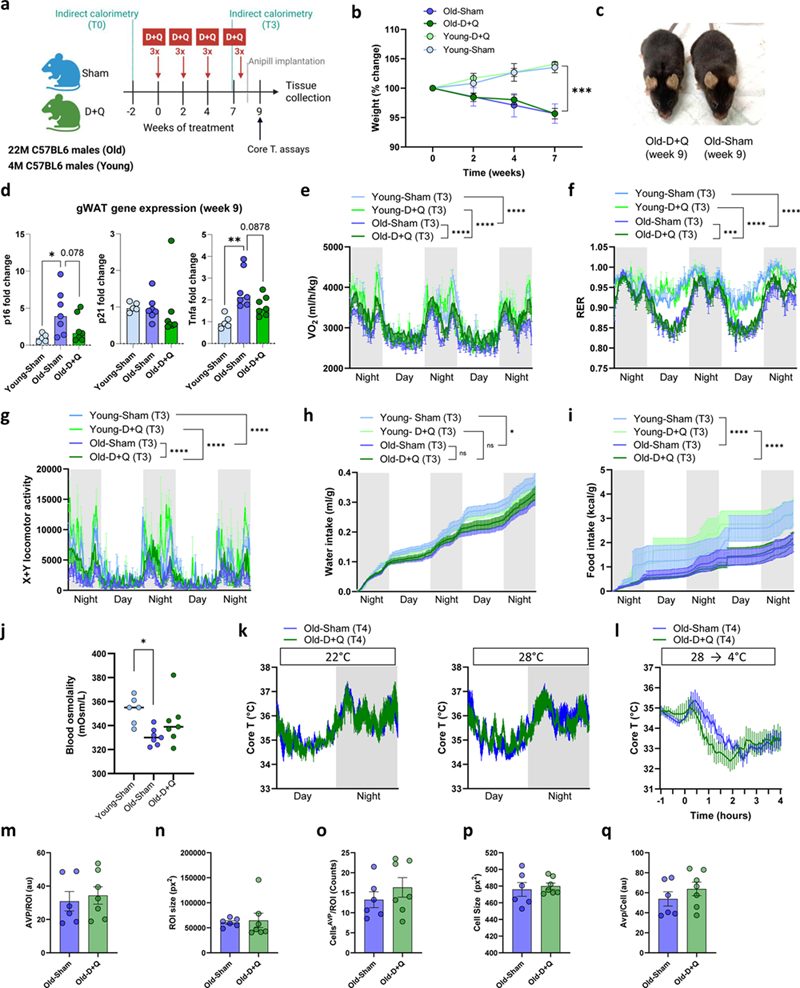
Senolytic treatment improves systemic metabolism but does not restore vasopressin-related defects in aged mice. **a.** Experimental design: Mice were divided into D+Q and Sham gavage experimental groups. **b.** Weight changes over time of old (*n = 7-8*) and young (*n = 3-4*). **c.** Representative images of D+Q and Sham-gavage mice. **d.** RT-qPCR of p16, p21 and Tnfa expression in gWAT biopsies after 9 weeks of treatment. **e-i.** Indirect calorimetry assessment of D+Q or Sham-treated mice measuring **(e)** VO_2_, **(f)** RER, **(g)** X+Y locomotor activity, **(h)** water intake, and **(i)** food intake. **j.** Osmolality derived from plasma, **k-l.** Core T measurements at 22°C and 28°C in the light and dark phases and **(k)** under 4°C challenge **(l)**. **m-q.** Quantification of morphological features of AVP^+^ neurons in each region: (**m)** AVP signal intensity per ROI, **(n)** ROI size, **(o)** AVP cell bodies per ROI, **(p)** AVP cell body size, and **(q)** AVP signal intensity per cell. Data are means ± sem. *One-way ANOVA or Student’s t-test (bar graphs), two-way ANOVA (line-graphs) were used. p-values: ns= p > 0.05; *** = p < 0.005; **** = p < 0.0005*.

## Discussion

Aging is accompanied by a progressive decline in multiple aspects of physiological homeostasis, including thermoregulation, metabolism, and fluid balance (**Fig. 1**). Our study identifies hypothalamic AVP neurons in the SON as a previously unrecognized driver of these shared age-related deficits. By combining single-nucleus transcriptomics, morphological and electrophysiological analyses, chemogenetics, and gene knockdown approaches, we reveal that SON^AVP^ neurons become transcriptionally and functionally hyperactive with age, leading to impaired water intake and altered temperature regulation.

We found that AVP was one of the most upregulated neuronal transcripts in the aged anterior hypothalamus, accompanied by soma enlargement and heightened excitability of SON^AVP^ neurons (**Fig. 2,3**). These cellular changes are consistent with enhanced peptide synthesis and release, as observed in both rodent and human studies of aging^37,40,41,67^. Functionally, acute chemogenetic activation of SON^AVP^ neurons in young mice reproduced hallmark features of aging, including hypothermia, reduced energy expenditure, and blunted water intake, underpinning the sufficiency of this circuit to drive age-like physiological decline (**Fig. 4**). Conversely, targeted knockdown of *Avp* in the SON of aged mice restored water consumption and modestly improved thermoregulatory control (**Fig. 6**), suggesting that excessive AVP signaling causally contributes to these phenotypes. Knockdown of *Avp* also improved carbohydrate metabolism in aged mice, while not altering body weight, adiposity or energy expenditure. These benefits could be attributed to potential effects on hepatic glucose output, pancreatic hormone secretion, or Hypothalamic Pituitary Adrenal (HPA) axis activity.

Water balance disruptions including hyponatremia and dehydration are more prevalent in older adults^11,68^. Thirst is regulated by salt-sensing CVO neurons including the subfornical organ (SFO) and organum vasculosum of the lamina terminalis (OVLT)^27,69^. However, aged mice retained their sensitivity to salt-induced thirst **(Fig. 1**) implying that age-associated dysfunctions in water balance are impaired by other neural components, possibly directly at the level of SON^AVP^ neurons. Increasing AVP drives lower blood sodium concentration by causing the kidneys to retain water, causing hyponatremia^70^. Our data suggest that high release of AVP during aging is causal for observed hyponatremia and reduced water intake, and that blood salt homeostasis is disrupted with age.

Concomitant with altered water balance, we also observed thermoregulatory dysfunction in aging. This defect is observed in older adults and is similarly associated with increased cognitive and health risks^71^. Remarkably, our chemogenetic results demonstrate that SON^AVP^ dysfunction might be a key driver of hypothermia and reduced energy expenditure with age (**Fig. 4**). Interestingly, our rescue experiments using *Avp* knockdown in the SON demonstrated a direct link between altering AVP levels and water balance but were less effective in alleviating age-related thermoregulatory deficits (**Fig. 6**). This dissociation suggests that while AVP is a critical hypothalamic output for maintaining water homeostasis, its contribution to thermoregulation may be more indirect or dependent on peripheral effector systems. It is well established that other organ systems are involved in controlling thermoregulation, including BAT-mediated thermogenesis and cutaneous vasoconstriction. Importantly, BAT undergoes profound age-related remodeling, including reductions in mass and cellular composition, leading to blunted thermogenic capacity^72^. White adipose tissue also becomes increasingly fibrotic and inflamed with age, further compromising metabolic flexibility and heat production^73^. These alterations have low plasticity once established and have so far been solely rescued by interventions at the level of adipose tissue^73^. Thus, even though SON-derived AVP signaling may influence thermoregulatory circuits, the age-related decline in adipose tissue function likely represents a key bottleneck that limits the ability of central AVP restoration alone to normalize thermal homeostasis. Our findings in mouse models support a critical role for low AVP levels in sustaining high energy expenditure during youth through modulation of BAT thermogenesis, whereas elevated AVP levels in aged mice are associated with reduced energy expenditure.

Pharmacological experiments further revealed dissociable roles for AVP receptor subtypes: V1A receptor signaling mediated thermoregulatory impairments, whereas both V1A and V2 receptors contributed to reduced water intake (**Fig. 5**). This receptor specificity highlights distinct mechanisms through which AVP exerts its effects on peripheral and central targets. Furthermore, pharmacological inhibition of V1A and V2 receptors in aged mice had similar results to that observed in chemogenetic mouse models for water intake benefits (**Extended Fig. 5**). Interestingly, the cell specific expression of V1AR receptor for Core T modulation is unclear as it is found on various organ systems, but its expression on vascular smooth muscle is likely to contribute to temperature regulation through its control of blood vessel vasoconstriction^74^. However, according to our data, BAT thermogenesis plays a dominant role over blood vessel dilation in driving hypothermia, as our two models of increasing plasma AVP yielded opposing effects on tail vasoconstriction (**Fig. 4 and Extended Data Fig. 5**). Corroborating our findings, V1A receptors are widely expressed in BAT and WAT tissues^75^.

Furthermore, we aimed at testing the impact of anti-aging therapies on our system using senolytic drugs. Although D+Q therapy improved systemic metabolic health and locomotor activity in aged mice, it did not significantly restore SON^AVP^ function or reduce hypothalamic inflammation, at least at the dose and timeframe examined (**Fig. 7, Extended Fig. 7**). This finding suggests that systemic “inflammaging”, which was alleviated by D+Q, might not contribute substantially to SON-specific neuronal hyperactivity with aging. However, as D+Q treatment did not reduce inflammatory IL-6 in the aged SON, we cannot rule out that accumulation of IL-6 or other inflammatory components in the SON could lead to dysfunction of AVP neurons, or that other senolytic agents with more complete penetration of the intact blood brain barrier, or more intense or longer treatment might be effective. It is plausible that IL-6 secreted within the SON regulates the activity of AVP neurons, as in humans, the intravenous delivery of IL-6 led to increased plasma AVP^76^.

The findings in this study position SON^AVP^ neurons as a critical hypothalamic hub linking the brain to physiological frailty by contributing to age-related impairments in water balance, thermoregulatory control and energy expenditure. These results broaden our understanding of the neural basis of homeostatic decline in aging and highlight AVP signaling as a potential therapeutic target. Interventions aimed at modulating hypothalamic AVP activity may therefore offer new opportunities to improve health span and resilience in aging. For example, interventions that normalize AVP signaling could alleviate chronic hypovolemia, a common age-associated condition characterized by reduced blood volume, which has been linked to brain volume loss, impaired cerebral microcirculation, altered synaptic function, glial activation, and synaptic loss^13–16^.

## Acknowledgements

We thank the Applied Genomics core at CSMC for their assistance in processing snRNA-seq. This work was supported by the Larry L Hillblom Foundation startup grant 2018-A-009-SUP (CER), the Cedars-Sinai Pilot Award from the Center on Aging and Diabetes (CER), the CIRM Scholar fund EDUC-12751 (NM), the National Institute of Aging grant RF1AG091203 (CER) and R37AG013925 (JLK, TT).

## Data and Code Availability

The corresponding author will provide the raw data supporting this manuscript’s findings on reasonable request.

## Material and Methods

### Animals

All procedures were approved by the Animal Care and Use Committee of Cedars Sinai Medical Center. In this study, we used C57BL/6 (Jax Strain 000664), C57BL/6 (NIA colony, Charles River), 5XFAD (Jax Strain 034848), and AVP-IRES2-Cre (Jax Strain 023530). Mice were bred in our colony according to Jackson Laboratory instructions for each strain. In all experiments, appropriate littermate controls were used. Mice were maintained on a 14 h:10 h light:dark cycle and fed normal chow (PicoLab Rodent 20 5053*, LabDiet).

### In vivo animal phenotyping and body composition

Indirect calorimetry (VO_2_, EE, RER), physical activity, water intake and food Intake data were collected in an automated home cage eight-chamber phenotyping system (Phenomaster v.6.6.1., TSE). Data sampling was every 18 minutes, with 2 minutes timeframe in each chamber. Body composition was assessed by EchoMRI. Individually housed mice were acclimated in the chambers for at least 24 hours. Food and water were provided *ad libitum* in the appropriate devices and measured by the built-in automated instruments. Locomotor activity and parameters of indirect calorimetry were measured for at least the following 72 hours. Anipill monitors for core temperature monitoring were surgically inserted into the abdominal cavity along the sagittal plane. Measurements were set to 5-minute intervals and collected for the duration of the mouse’s lifespan. Interscapular BAT and tail temperatures were obtained with Flir E60bx or E75 thermal cameras and analyzed with QuickReport software to extract temperature recordings of the BAT area and the base of the tail. Osmotic thirst was induced by an intraperitoneal injection of 2M NaCl (5 μl/g bodyweight).

### SnRNA-seq of the anterior hypothalamus

The anterior hypothalamus region containing the POA (Bregma A/P +0.5 to −0.6 mm) were dissected together from 5 young wild-type male mice (Age 3m) and 5 aged mice (Age 22m). Dissected tissues from 5 mice were pooled for each sample, dounce homogenized on ice and centrifuged at 500g for 10 min at 4 °C. Pellets were resuspended and spun in an iodixanol gradient at 3000g for 1hr, 4 °C. Myelin was removed using Myelin Removal Beads II (Miltenyi Biotec, Cat#130-096-733) using LS columns (Miltenyi Biotec, Cat#130-042-401) according to the manufacturer’s instructions and the extracted cell layer was resuspended in 1% BSA solution with RNAse inhibitor. Nuclei were pelleted at 500g for 10 minutes at 4 °C before final resuspension. The concentration of nuclei was adjusted as per 10x chromium chip requirements. Supernatants were removed and passed through a 70-μm filter then through a 20-μm filter (MACS Smartstrainer) to remove large debris. DAPI was added to the suspensions and nuclei were separated by FACS. The resulting suspensions were loaded into a 10x Genomics Chromium single-cell chip at a concentration of about 700-1200 nuclei per μl with the aim of sequencing 10000 nuclei per sample. Downstream preparation of sequencing libraries was carried out using the 10x Genomics Single Cell Kit V2. The libraries were sequenced on an Illumina NextSeq500 instrument using instructions provided by 10x Genomics. Paired-end sequencing with read lengths of 150 nt was performed for all samples. Illumina sequencing reads were aligned to the mouse genome using the 10x Genomics CellRanger pipeline with the default parameters.

### Seurat workflow and neural analysis

Identification of cell-type clusters was performed as previously described using the Seurat package in R. In brief, we first filtered out non-nuclei such as mitochondria and red blood cells as well as obvious doublets. Data were scaled by Seurat::ScaleData then underwent linear dimensional reduction. To achieve this, we identified variable features by the variance stabilizing transformation (Vst) selection method and used those features in PCA dimension reduction. To generate cell clusters, we computed nearest neighbors clustered cells by a shared nearest neighbor (SNN) modularity optimization based clustering algorithm using the Seurat commands FindNeighbors (dimensionality reduction of 40) and FindClusters (resolution set to 0.9), respectively. The resulting clusters were visualized by 2D UMAP. We used the following markers to identify major cell type classes: neurons (Eno2+, Snap25+, Tubb3+, Ndrg4+, Stmn2+), microglia (Slco1c1+,Fn1+,Slco1a4+,Cldn5+), astrocytes (Gfap+,Ntrs2+,Ntsr2+,Aldoc+), mature oligodendrocytes (Mag+,Mbp+,Utd8+,Mog+), oligodendrocyte precursors (Cspg4+,Pdgfra+), tanycytes (Rax+), fibroblasts (Col1a1+) and endothelial cells (Slco1a4+,Cldn5+,Fn1+,Slco1a4+). Neural populations were subset using excitatory (Slc17a6+) and inhibitory (Slc32a1+) markers and underwent further normalization by methods previously described and visualized using UMAP. Genes of interest were investigated across new clusters via Seurat::DotPlot.

### RNA velocity analysis of glial cells

Datasets were reprocessed and clustered using Scanpy in Python. Cell-types were subset and reclustered before performing velocity and pseudotime analysis.

### Stereotaxic viral injections

Anesthesia was induced using isoflurane (induction, 5%; maintenance, 2–3%) and administered Carprofen (5 mg/kg body weight) and buprenorphine (0.1 mg/kg body weight) before surgical procedure subcutaneously (sc). Lidocaine (5mg/kg) was injected sc prior to making a cranial incision. A 0.5mm hole was drilled into the skull over the target site. All injections were done using a microliter syringe (Hamilton) and UMP3 pump (WPI) mounted directly on the stereotactic frame and the viruses were delivered at a rate of 2 nl sec-1. Injections were administered bilaterally (one at a time) in the SON (coordinates relative to bregma: A/P −0.6 mm, M/L ±1.10 mm and D/V −5.4 mm). After the injection, the needle of the syringe was kept in place for 10 min to reduce backflow and then withdrawn slowly. Animals recovered on a heating pad until normal behavior resumed and monitored. Additional doses of carprofen were administered 24- and 48-hours post-surgery, while buprenorphine was given every 8-12 hours for 48 hours post-surgery.

### In vivo chemogenetics and AVP peptide assays

For chemogenetic experiments, 150 nl of AAV-hSyn-DIO-hM3D(Gq)-mCherry (Addgene, 44361-AAV8) was injected on each side. After two-week recovery, animals were acclimatized for the first 2 days with saline (i.p) and on day 3, mice received treatment. All injections were performed at the same time (12pm) Mice that had received viral vector were administered DCZ (Tocris, 7193) at 0.1 or 1mg/kg body weight. For AVP peptide experiment, mice were acclimatized for the first 2 days with saline (i.p) and on day 3, mice received AVP (ToCris, 2935) at 0.1mg/kg. All injections were performed at the same time (12pm). For the antagonist experiments, animals were acclimated for two days with saline (i.p) followed by treatment. For antagonists, mice received I.P. injection of antagonist or saline 20 minutes prior to stimulus. All antagonist experiments were performed at the same time (t=0 as 6pm) to decipher changes between experiments. The following antagonists (10mg/kg) were used for V1AR, V2R, and V1BR, respectively: SR 49059 (Tocris, 2310), Tolvaptan (Sigma, T7455), and TASP 0390325 (Tocris, 5729).

### *Avp* Genetic silencing

Mice were injected on each side 0.5uL of lentiviral particles encoding either SMARTvector Lentiviral Mouse *Avp* mCMV-TurboGFP shRNA, 10^8 TU/mL (Horizon) or SMARTvector Non-targeting mCMV-TurboGFP Control Particles, 10^8 TU/mL (Horizon).

### D+Q senolytic cocktail

Dasatinib (SML2589, Sigma-Aldrich) and Quercetin (Q4951, Sigma-Aldrich) were reconstituted in vehicle (10% ethanol, 30% polyethylene glycol 400, 60% Phosal 50 PG) solution. Drug cocktail was delivered via oral gavage at 5mg/kg Dasatinib and 50mg/kg Quercetin dose. Three doses were delivered at approximately the same time (4pm) every other day for each round of treatment.

### Tissue Harvest and Processing

In consideration of circadian oscillations, all mouse endpoints were performed between 12-4pm. Mice were anesthetized with isoflurane and blood was drawn by cardiac puncture into heparin-coated tubes (BD vacutainer). This was followed by perfusion with ice-cold 0.1M potassium phosphate-buffered saline (PBS, pH 7.4). Brains were removed, stored in 4% paraformaldehyde (PFA), and transferred into 30% sucrose at 4°C for 48 hours. The PFA-fixed brains were embedded in optimal cutting temperature (OCT) compound (Tissue-Tek) and cut into 40 microns coronal sections of the SON (Bregma −0.45 to −0.8), PVN (Bregma −0.55 to −1.0), and SCN (Bregma −0.18 to −0.8) on a Leica cryostat.

### Plasma Assays

Blood collected in heparin coated tubes (BD vacutainer) sat in ice for approximately 30 minutes, then spun at 2000 x g for 15 minutes to extract plasma. Osmolality measurements were conducted within the Pathology and Lab Medicine Core using OsmoPRO® Multi-Sample Micro-Osmometer (Advanced Instruments). AVP levels and copeptin levels were measured using the commercially available Arg8-Vasopressin Competitive ELISA (Invitrogen, EIAAVP) or Mouse Copeptin ELISA (Novus Biologicals, NBP3-42239) kits.

### Fluorescent in-situ hybridization

In-situ hybridization was done using RNAscope® multiplex fluorescent reagent kit v2 (323100, ACD Bio), following the manufacturer’s protocols. Avp was detected by RNAscope® probe Mm-Avp (401391, ACD Bio). Slides were cover slipped with ProLong™ Glass Antifade Mountant (P36980, Fisher Scientific).

### Immunohistochemistry

Brain sections were incubated in PBS-TritonX blocking solution containing 2% normal donkey serum and 0.2% Triton X-100 for 1 hour at room temperature, followed by primary antibody overnight. The sections were washed three times in PBS with Tween20 (0.5%) and then incubated in one of the following secondary antibodies for 1h in the dark at room temperature. Sections were washed three times in PBST, mounted onto slides, and stained with either Nissl NeuroTrace™ 640/660 (N21483, Invitrogen) and cover slipped with ProLong™ Glass Antifade Mountant (P36980, Fisher Scientific). Alternatively, they were counterstained and coverslipped in DAPI stain mountant (P36931, Life Technologies). The following antibodies were used: rabbit anti-AVP (1:1000, ab213708, Abcam), rat anti-IL-6 (1:250, AMC0864, Invitrogen), rabbit anti-GFAP (1:500, PA1-10019, Invitrogen), anti-NeuN (1:1000, ab279297, Abcam), anti-cFos (1:1000, ab190289, Abcam) and anti-mCherry (1:500, PA5-34974, Thermo Fisher Scientific), The sections were washed three times in PBS with Tween20 (0.5%) and then incubated in one of the following secondary antibodies for 1 hour in the dark at room temperature: donkey anti-rabbit Alexa Fluor® 488(1:600, A21206, ThermoFisher) and donkey anti-rat Alexa Fluor® 488 (1:600, A-21208, ThermoFisher). Sections were washed three times in PBST and mounted onto slides coverslipped with ProLong™ Glass Antifade Mountant (P36980, Fisher Scientific).

### Imaging and Analysis

Fluorescent sections of SON, PVN and SCN were imaged with BZ-X700 Keyence at 20x magnification. Higher magnification imaging (63-100x) was accomplished via Apotome Zeiss Axiocam. To perform qualitative and quantitative analysis of Avp cell bodies in SON, PVN and SCN six sections containing both hemisheres for each region of interest (ROI) were imaged at 20x then annotated by a blinded researcher using Qupath (v 0.3.2). Area (px^2) of ROI were based on these annotations. Avp signal intensities were calculated for the indicated ROI [*Avp/ROI*] with a resolution downsample of 1 and tile diameter 200 px. Cell counts per ROI were generated by native ‘Cell Count’ function in Qupath (sigma set to 2, minimum area 100px^2, threshold 25). Shape feature values were generated including [*<u>area (px^2)/cell</u>*] and [*circularity/cell*]. Intensity features were also generated for individual detections to determine [*Avp/Cell*]. Measurements from ROI and cell-based metrics were then streamlined and averaged across each sample in R. To count detections of IL-6 puncta and GFAP+ cell bodies, images were processed using Macro Cell Count in BZ-X Analyzer Software.

### Ex vivo electrophysiology - Slice preparation

Brains from 5 young mice aged between 3-5-month-old and 5 old mice, aged between 15–16-month-old were used. Animals were deeply anaesthetized with ketamine and xylazine, until no response to a tail pinch was observed. Then, they were perfused transcardially with ice-cold, carbogenated (95% O2, 5% CO2) cutting artificial cerebro-spinal fluid (cACSF), composed of (in mM): 75 NaCl, 2.5 KCl, 6 MgCl2, 1.2 NAH2PO4, 0.1 CaCl2, 25 NaHCO3, 2.5 glucose, 64 sucrose, 5 sodium ascorbate, 3 sodium pyruvate and 2 thiourea. This was followed by decapitation, brain extraction and slicing into 250 μm thick coronal slices with a vibroslicer (Leica VT1000S, Leica Biosystems). Slices containing SON were transferred to an incubation chamber with RT cACSF for about 30 mins, after which they were placed in carbogenated normal ACSF (nACSF; warmed up to 32C), which contained (in mM): 115 NaCl, 2.5 KCl, 1.2 MgCl2, 1.2 NaH2PO4, 2.5 CaCl2, 21.4 NaHCO3, and 11.1 glucose.

### Patch clamp recordings and immunohistochemical verification

After a minimum of 2h of incubation, brain slices were placed in a recording chamber, perfused with carbogenated, heated to 32C nACSF at a speed of 1.5 ml/min. nACSF was suplemented with NMDA, AMPA, GABAA and GABAB receptor blockers (10 μM MK801, 10 μM DNQX, 100 μM picrotoxin, 0.3 μM CGP55845, respectively), in order to exclude potential network modulation by various glutamate and GABAergic inputs and record cells’ spontaneous activity. Recording chamber was positioned under an Olympus BX31WI microscope fitted with infrared differential interference contrast. Whole-cell configuration was acquired under 40x magnification with borosilicate glass pipettes (TW150F-4, WPI; resistance = 5–10 MΩ), filled with an intrapipette solution containing (in mM): 110 potassium gluconate, 10 HEPES, 10 KCl, 2 MgCl2, 0.5 Na-ATP and 0.05% biocytin (pH = 7.4, adjusted with KOH; osmolality 300 mOsmol/kg adjusted with sucrose) and mounted onto an Ag/AgCl electrode. All recordings were made in current clamp mode, using Clampex software (version 10.7.0.3, Molecular Devices). Recorded signal was low-pass filtered at 10 kHz and recorded at 20 kHz with Multiclamp 700B (Molecular Devices. The seal test was used to determine the values of membrane capacitance and resistance upon establishing whole-cell configuration. The resting membrane potential and spontaneous firing rate were recorded at a holding current of 0 nA for at least 30s. Following, each cell was manually brought down to −65 mV (below firing threshold) by negative current injection, from which a 1 s 0.5 nA current ramp was applied in order to determine the threshold for action potential (AP) generation and highest possible firing rate before entering depolarization block. Lastly, a set of depolarizing current steps ranging from 0 to 100 pA (10 steps with 10 pA increments, 0.5s long) were applied, to determine the gain in firing rate with increasing stimulus strength. A liquid junction potential of −15 mV was added to the recorded values. All brain slices containing the recorded cells were immunohistochemically stained to confirm their location within the SON and verify whether they contained AVP. Immediately after recording, the slices were fixed overnight in 4% paraformaldehyde in phosphate-buffered saline (PBS, pH = 7.4). On the following day, PFA was washed out with PBS (3 x 5 min) and the slices underwent membrane permeabilization and non-specific site blocking with 0.2% Triton-X100 and 4% normal goat serum (NGS) in PBS, for 1 h at room temperature. Next, the slices were incubated with primary rabbit AVP antibody and processed for secondary antibody as described in immunohistochemistry section. To visualize recorded neurons, this solution also contained secondary antibodies Alexa647-conjugated Streptavidin (1:1000, Invitrogen #S21374), which binds biocytin present in the intrapipette solution. At the end, after another 3×5 min wash in PBS slices were placed onto microscope slides and coverslipped with Fluoromount-G® (SouthernBiotech). Images were taken with a confocal microscope (Nikon AX R NSPARC, Nikon Instruments Inc.).

### Quantitative PCR analysis of adipose tissue

RNA was isolated using TRIzol/chloroform extraction and RNEasy Plus Micro Kit (Qiagen, 74034). Subsequently, 1 µg of RNA was converted to cDNA using the M-MLV Reverse Transcriptase kit (Invitrogen). Quantitative PCR was performed on a QuantStudio 7 Pro instrument with PerfeCTa qPCR FastMix II (QuantaBio). Gene expression levels were normalized to TATA-binding protein (TBP) and analyzed by the 2-ΔΔCT method using Design & Analysis Software v2.8.0. Mouse TaqMan probes from Mm.pt.58.42804808 (p16Ink4a, Integrated DNA Technologies), Mm04205640_g1 (p21cip1, Thermo Fisher Scientific), Mm00443258_m1 (tnfa, Thermo Fisher Scientific).

### Statistical analysis

GraphPad Prism 9 was used for statistical analysis. Data are expressed as mean ± SEM. p Values were calculated using Student t-test, one-way ANOVA, two-way ANOVA or mixed models followed with appropriate post *hoc* tests. Analysis of covariance (ANCOVA) was performed in R. p values less than 0.05 were considered significant.

## Notes

### Competing Interest Statement

The authors have declared no competing interest.

