## supplemental data for "The Role of Supraoptic Hypothalamic Arginine Vasopressin Neurons in Aging-Associated Water Balance and Thermoregulatory Deficits"

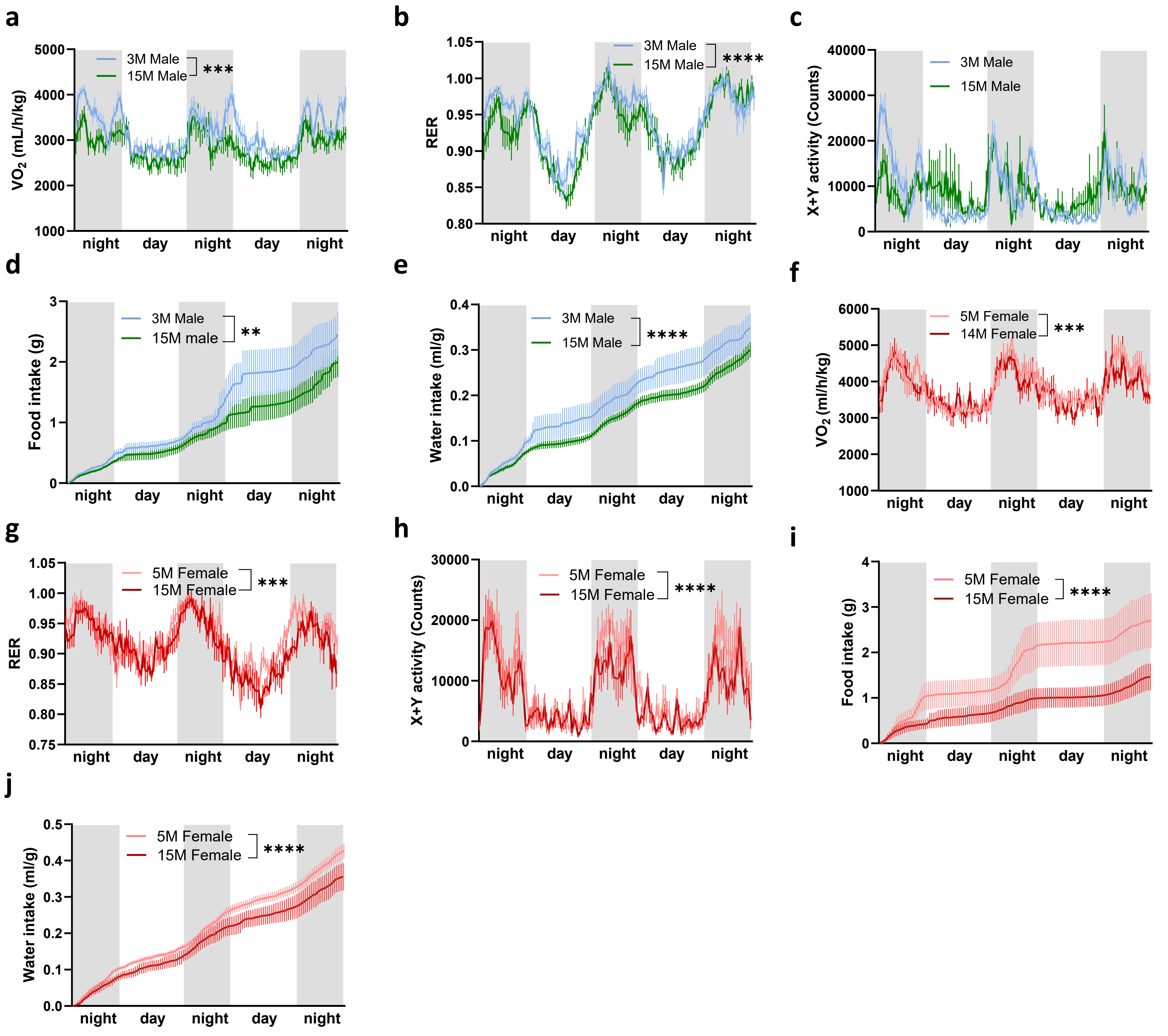


**Extended Data Fig. 1. Metabolic changes in middle-aged male and female mice.** Indirect calorimetry of male (**a-e**) and female (**f-j**) mice by age over three days: **a,f**. $VO_2$, **b,g.** RER, **c,h.** locomotor activity, **d, i.** food intake, and **e, j.** water intake. Data are means ± sem. *Two-way ANOVA. p-values: ns = p > 0.05; *** = p < 0.005; **** = p < 0.0005.*

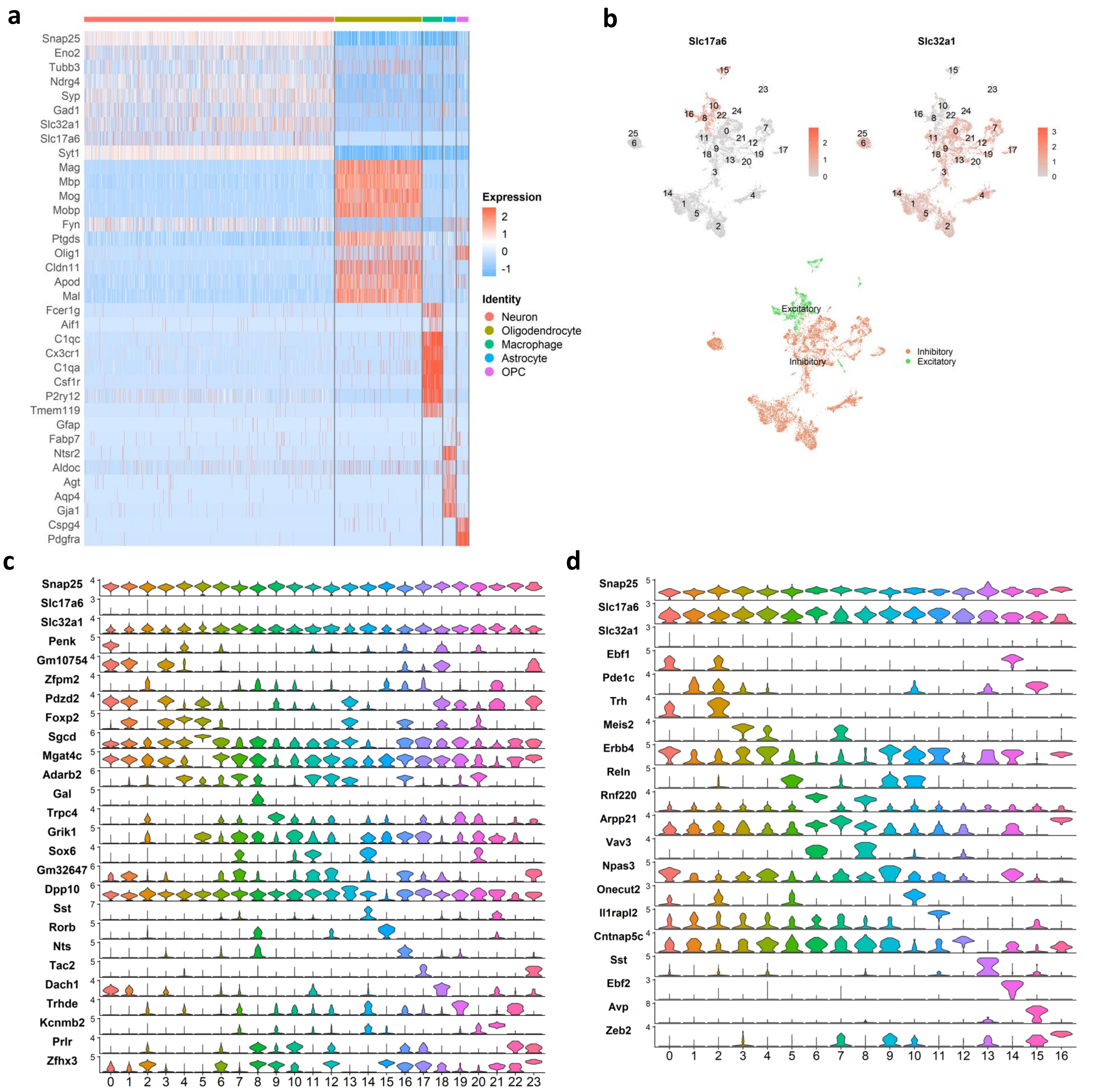


**Extended Data Fig. 2. Single-nuclei RNA-seq analysis of neural cell-types. a.** Heatmap of cell-specific markers show consistency in cell identity. **b.** The excitatory neural marker Slc17a6 and inhibitory neural marker Slc32a1 are shown to distinguish neural subtypes. **c.** Inhibitory and **d.** excitatory clusters were subclustered from the neuronal pool. Most expressed genes in each subcluster were identified by FindAllMarkers and shown in violin plots.

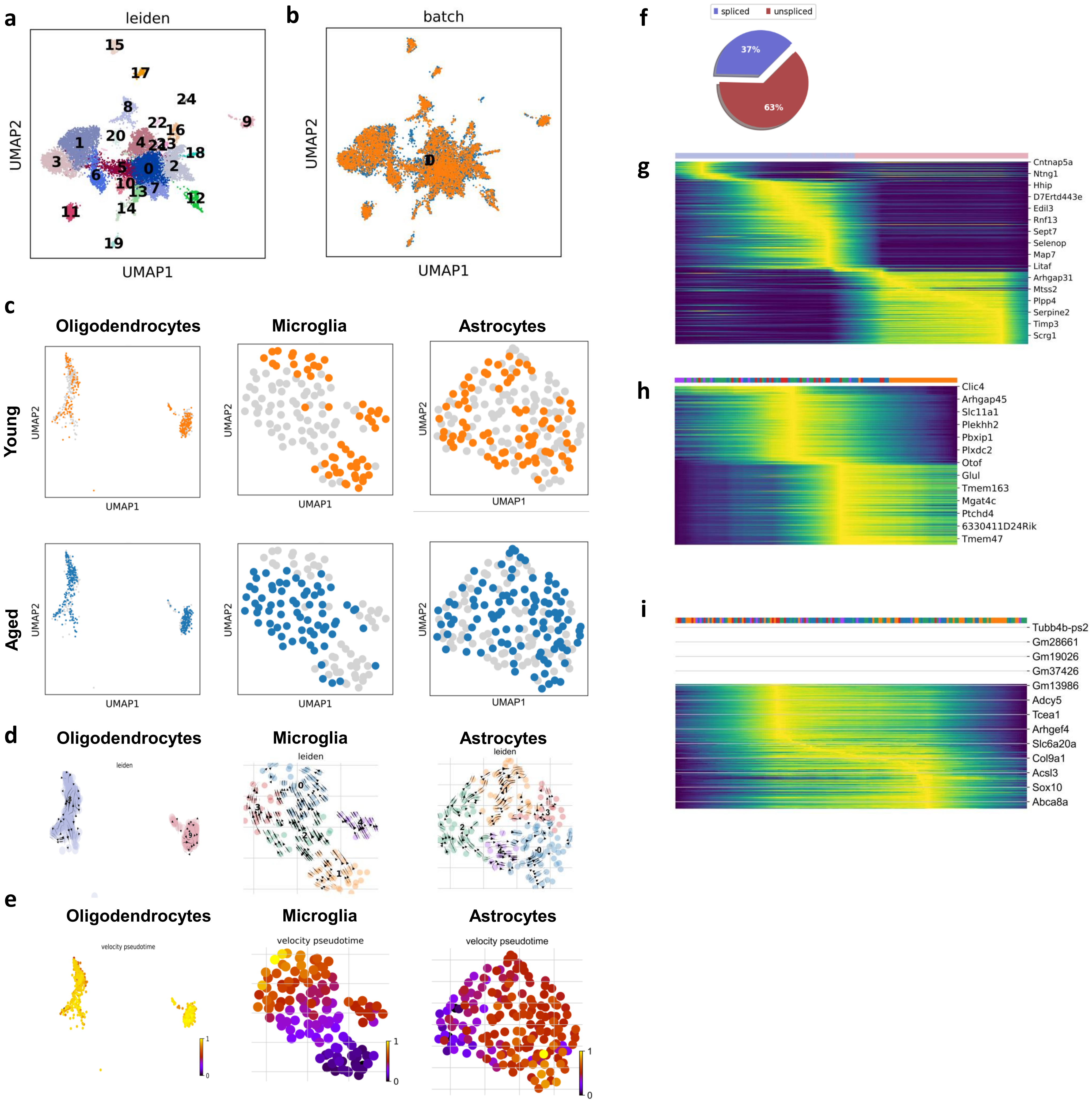


**Extended Data Fig. 3. RNA Velocity Analysis of glial cells. a-b.** UMAP of (a) cell clusters and (b) superimposed young and aged sample pools. **c.** Cells labeled by group for subclustered oligodendrocyte, microglia and astrocyte. **d.** Directionality of cell-specific subclusters. **e.** RNA velocity pseudotime plotted. **f.** RNA spliced vs. unspliced proportions of oligodendrocytes. **g-i.** Heatmaps of RNA driver genes for **g.** oligodendrocyte, **h.** microglia, and **i.** astrocytes.

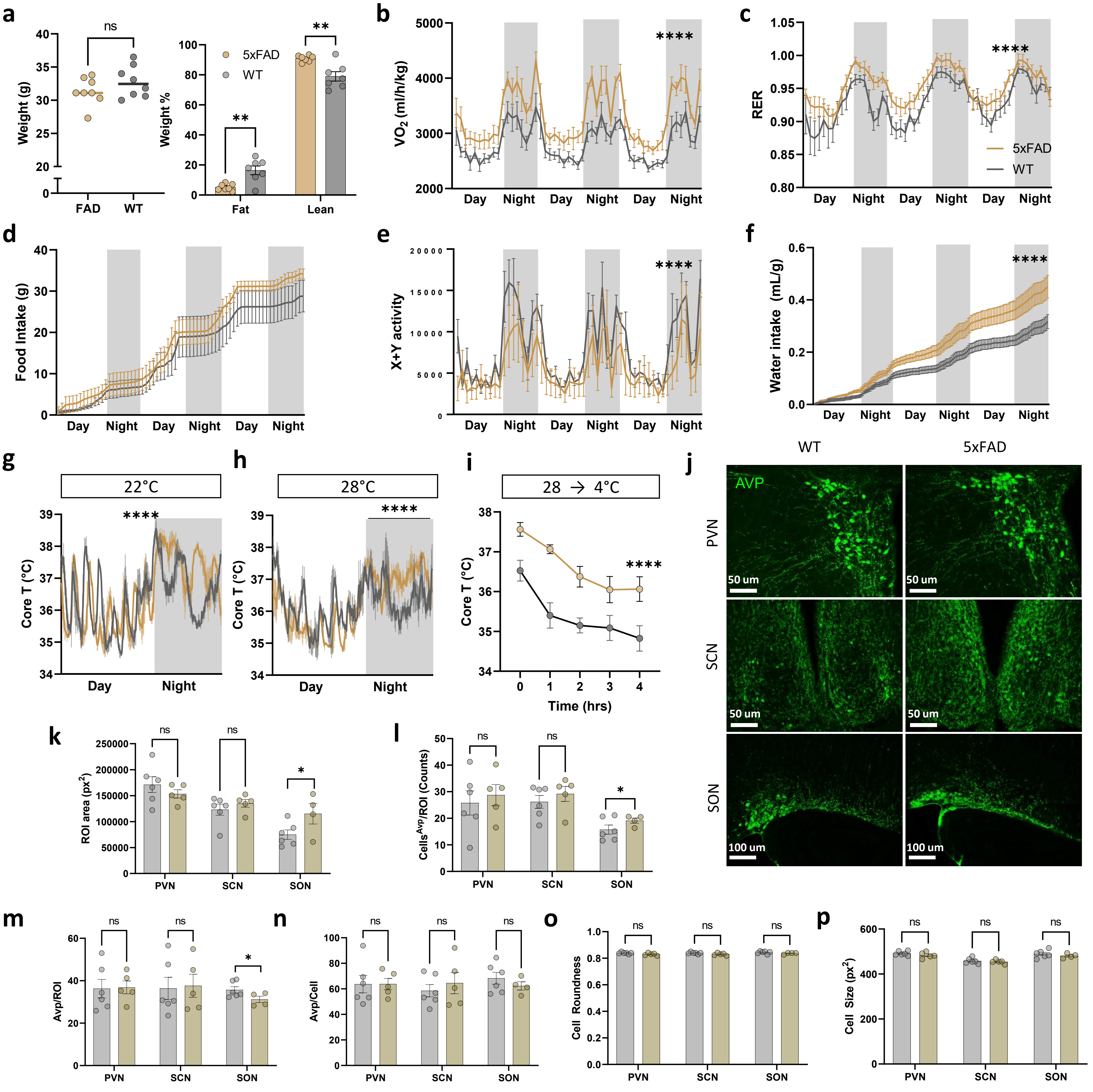


**Extended Data Fig. 4. Metabolic and thermoregulatory changes in 5xFAD mice. a.** Body weight (left) and body composition (right) of 6-month-old male 5XFAD and WT littermates, n=8. **b-f.** Indirect calorimetry over three days: (**b**) $VO_2$, (**c**) RER, (**d**) food intake, (**e**) locomotor activity and (**f**) water intake. **g.** Body core temperature (Core T) readings using intraperitoneal Anipill probes over 1 day and 1 night cycle (*n* young = 4, *n* aged = 3) at (**g**) 22 °C and (**h**) 28 °C environmental temperatures. **i.** Core T readings using a rectal probe during an acute cold challenge at 4 °C (*n* = 8). **j.** Representative immunofluorescence images of PVN, SCN and SON in 5xFAD and WT littermates. Qualitative measurements for ROI size (**k**), AVP cell bodies per ROI (**l**), AVP signal intensity per ROI (**m**), AVP cell body size (**n**), AVP cell roundness (**o**), and AVP signal intensity per cell (**p**). Data are means ± sem. *Student's t-test (a, k-p) or two-way ANOVA (b-i). p-values: ns = p > 0.05; *** = p < 0.005; **** = p < 0.0005.*

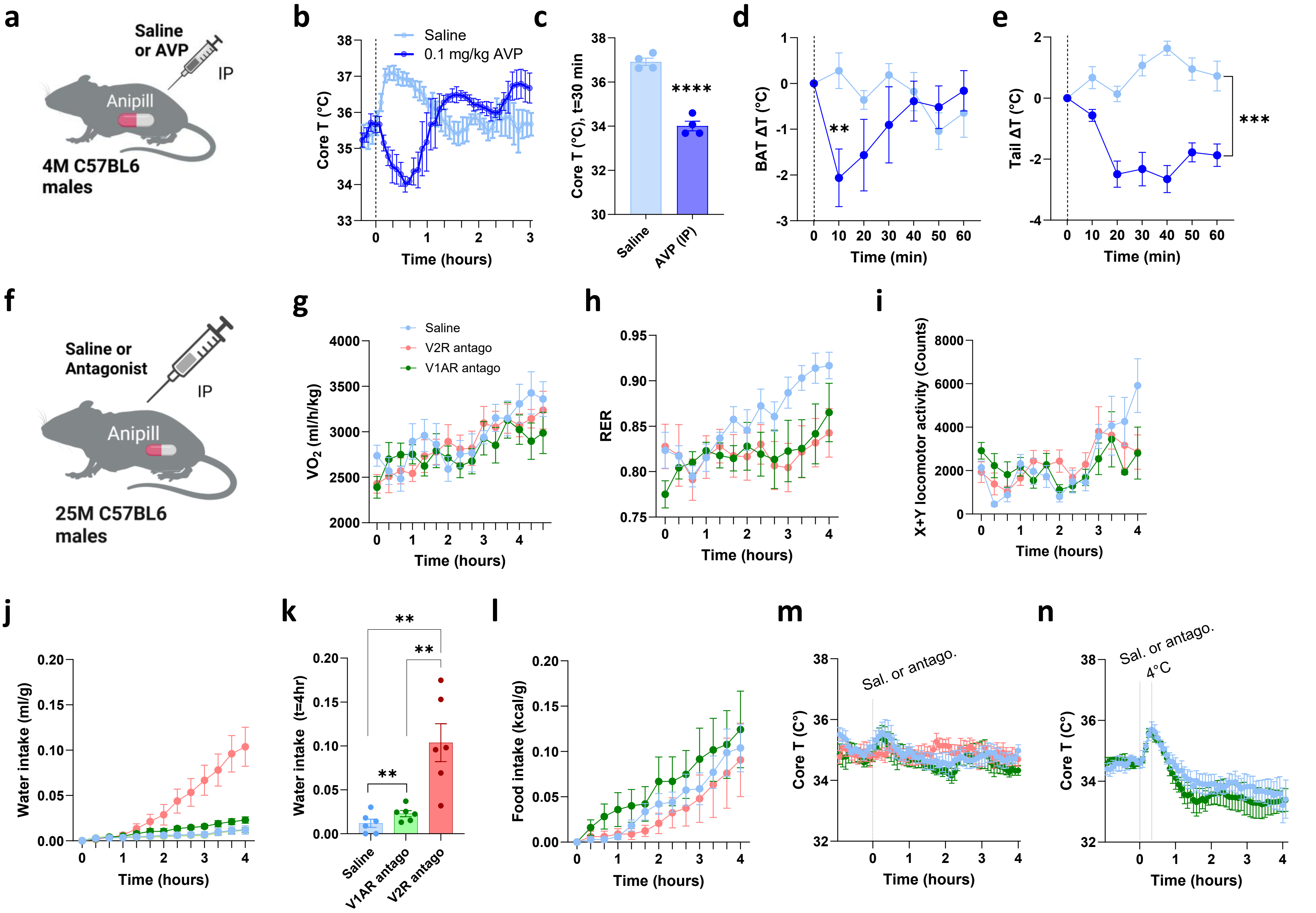


**Extended Data Fig. 5. Effects of peripheral vasopressin on physiological frailty with age. a.** Experimental design for (**b-e**): C57BL6/J mice with Anipill monitors in abdominal cavity were injected I.P. with either saline or 0.1mg/kg AVP peptide. Core T over time (**b**) and at t=30 min post-injection (**c**). **d-e.** Infrared (IR) temperatures delta changes from initial measures of BAT and Tail base. **f.** Experimental design: 25-month (25M) C57BL6 mice were injected (I.P.) with saline or antagonist. **g-i.** Indirect calorimetry assessment of vasopressin receptor antagonists: (**g**) $VO_2$, (**h**) RER, (**i**) locomotor activity, (**j**) water intake and (**l**) food intake. **m,n.** Core T readings post-injection at (**m**) 22 °C and (**n**) prior to starting 4°C cold challenge from 28°C. Data are means ± sem. *Student's t-test or two-way ANOVA. p-values: ns = p > 0.05; *** = p < 0.005; **** = p < 0.0005.*

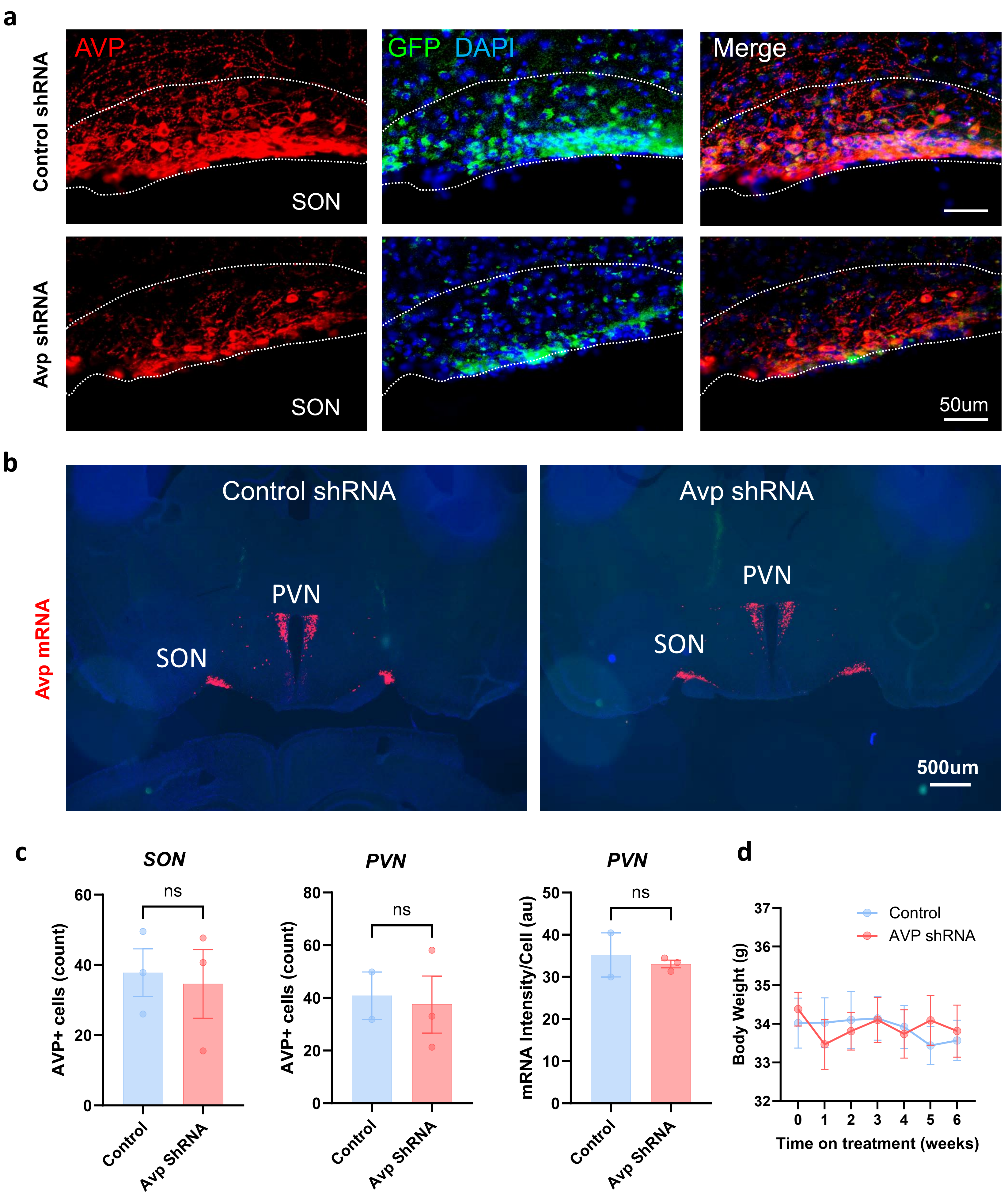


**Extended Data Fig. 6. Validation and qualitative analysis of SON$^{AVP}$ silencing. a**. Representative immunofluorescence SON images of control and AVP shRNA (SON$^{AVP}$) treated mice. **b.** RNAscope images of Avp expression within the aHYP region in control and shRNA treated mice. **c.** quantification of Avp+ cells in SON and PVN as well as Avp mRNA signal intensity in the PVN. **d.** Raw values of body weights post-treatment with lentiviral particles. Data are means ± sem. *Student's t-test or two-way ANOVA. p-values: ns = p > 0.05; *** = p < 0.005; **** = p < 0.0005.*

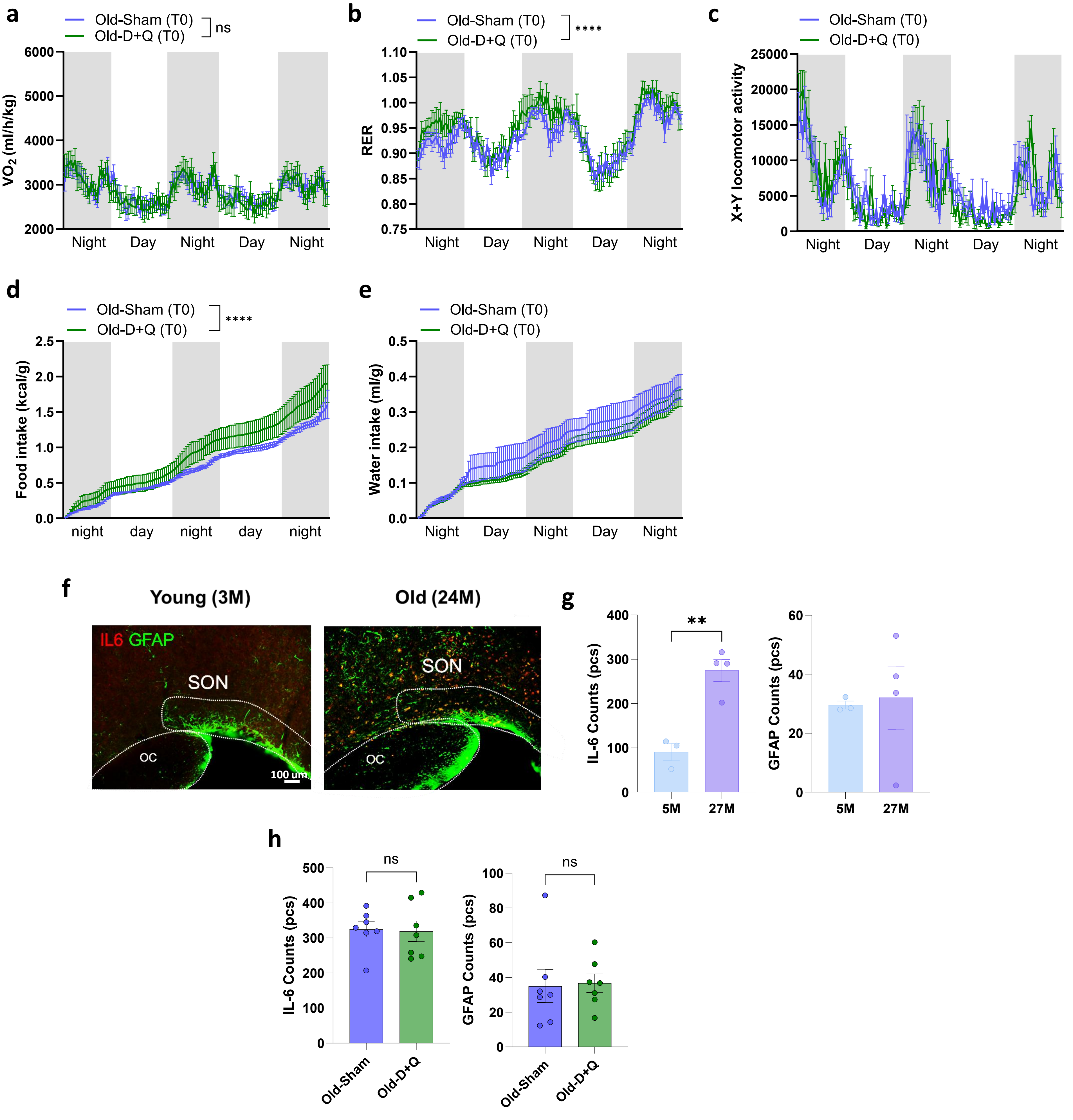


**Extended Data Fig. 7. Assessment of metabolic and inflammatory signals in D+Q treated mice. a-e.** Indirect calorimetry assessment of Old-Sham and Old-D+Q at t=0, or prior to receiving treatment, (**a**) $VO_2$, (**b**) RER, (**c**) X+Y locomotor activity, (**d**) food and (**e**) water intake over time. **f.** Representative image of immunofluorescence of SON used for qualitative analysis, anti-IL-6 (red), anti-GFAP (green). **g.** Quantification of immunohistochemical fluorescence counts for IL-6 or GFAP in 5-month (5M) vs. 27-month (27M) mice. **h.** Quantification of immunohistochemical fluorescence counts for IL-6 or GFAP in Old-Sham vs. Old-D+Q mice. Data are means ± sem. *Student's t-test or two-way ANOVA. p-values: p-values: ns = p > 0.05; *** = p < 0.005; **** = p < 0.0005.*
